# A versatile bacterial innate immunity protein directly senses two disparate phage proteins

**DOI:** 10.1101/2024.05.10.593582

**Authors:** Tong Zhang, Albinas Cepauskas, Anastasiia Nadieina, Aurelien Thureau, Kyo Coppieters ’t Wallant, Chloé Martens, Daniel C. Lim, Abel Garcia-Pino, Michael T. Laub

## Abstract

Eukaryotic innate immune systems use pattern recognition receptors (PRRs) to sense infection by detecting pathogen-associated molecular patterns, which then triggers an immune response. Bacteria have similarly evolved immunity proteins that sense certain components of their viral predators known as bacteriophages^1–6^. Although different immunity proteins can recognize different phage-encoded triggers, individual bacterial immunity proteins have only been found to sense a single trigger during infection, suggesting a one-to-one relationship between bacterial PRRs and their ligands^7–11^. Here, we demonstrate that the anti-phage defense protein CapRel^SJ46^ in *Escherichia coli* can directly bind and sense two completely unrelated and structurally different proteins using the same sensory domain, with overlapping but distinct interfaces. Our results highlight the remarkable versatility of an immune sensory domain, which may be a common property of anti-phage defense systems and enable them to keep pace with their rapidly evolving viral predators. We found that Bas11 phages harbor both trigger proteins that are sensed by CapRel^SJ46^ during infection, and we demonstrate that such phage can only fully evade CapRel^SJ46^ defense when both triggers are mutated. Our work reveals how a bacterial immune system that senses more than one trigger can help prevent phages from easily escaping detection, and it may allow detection of a broader range of phages. More generally, our findings illustrate unexpected multifactorial sensing by bacterial defense systems and complex coevolutionary relationships between them and their phage-encoded triggers.

## Introduction

A central facet of innate immunity is the use of pattern recognition receptors (PRRs) that bind specific pathogen-associated molecular patterns (PAMPs), leading to the activation of cell-intrinsic defense mechanisms^12,13^. In mammals, there are a diverse set of PRRs that recognize different PAMPs, *e.g.* RIG-I binds dsRNA, TLR4 binds LPS, and TLR5 binds flagellin^13^, with most mammalian PRRs specific for a single PAMP. There is at least one example, human NAIP/NLRC4, that can recognize three different ligands – bacterial flagellin, and the needle and inner rod proteins of the type III secretion system, but a common structural motif is recognized in each protein^14–16^.

The concept of PRRs and PAMPs extends to bacteria and their ability to detect infection by bacteriophages. However, the function, specificity, and evolution of innate immunity proteins in bacteria remain poorly understood. Recent work has indicated that bacteria often harbor proteins analogous to PRRs that recognize certain phage proteins or nucleic acids following infection, leading to the activation of various anti-phage defense mechanisms^7–11,17^. There are, as yet, only a handful of cases for which the direct trigger of an anti-phage defense system is known, so the specificity of phage detection by bacterial PRRs is largely unknown. Prior work has demonstrated that homologs of a given family of PRRs sometimes recognize different PAMPs^8^, and a large-scale screen indicated that some bacterial defense proteins can be activated by the ectopic expression of multiple unrelated phage proteins^17^. However, individual defense systems have only been reported to recognize single ligands during a phage infection suggesting that bacterial PRRs have one-to-one relationships with their activating phage-derived triggers.

Here, we demonstrate that the anti-phage defense protein, CapRel^SJ46^, can directly bind and sense two completely unrelated phage proteins using the same sensor domain. This multi-factorial sensing capability enables the defense system to recognize different phages that only produce one trigger protein or the other. Additionally, we found one phage that harbors both trigger proteins and we demonstrate that they both activate CapRel^SJ46^. As a result, CapRel^SJ46^ provides stronger defense against this phage compared to phages harboring either trigger protein alone. The ability to sense and respond to multiple trigger proteins may be a common and highly advantageous property of bacterial anti-phage defense systems as it can provide protection against a broader set of phages and, in some cases, make it more difficult for phages to acquire a single mutation that enables complete defense evasion. Further, our results indicate that the Red Queen dynamic unfolding between bacteria and phages may not always involve a single PRR and a single ligand, but rather involve the complex coevolution of multiple factors.

### Gp54^Bas11^ is an alternative activator of CapRel^SJ46^

We recently identified and characterized a fused toxin-antitoxin system called CapRel^SJ46^ that protects *E. coli* against diverse phages^11^. CapRel^SJ46^ contains an N-terminal toxin domain and a C-terminal antitoxin domain that normally binds and auto-inhibits the N-terminal toxin. During infection by SECΦ27 phage, the newly synthesized major capsid protein (MCP) binds directly to the C-terminal domain of CapRel^SJ46^ to relieve autoinhibition, leading to the activation of CapRel^SJ46^. Activated CapRel^SJ46^ then pyrophosphorylates the 3’ end of tRNAs, which inhibits protein translation and restricts phage propagation^11^. Sensing the major capsid protein, which is an essential and abundant component of the phage, is beneficial to the host bacteria because it limits the number of mutations that phages can acquire to escape defense. However, given intense selective pressure to maintain infectivity, phages can evolve to overcome defense through mutations in its capsid protein. For instance, a SECΦ27-like phage called Bas4 naturally encodes a single amino acid substitution in its MCP that prevents activation of CapRel^SJ46^, enabling the phage to escape defense^11^. Such escape may drive selection for mutations in CapRel^SJ46^ that restore an interaction with the MCP. Alternatively, CapRel^SJ46^ could, in principle, evolve to sense a different phage protein.

To explore whether CapRel^SJ46^ can sense phage factors other than the major capsid protein, we focused on a family of phages from the BASEL collection^18^ that are closely related to SECΦ27, including Bas11. When CapRel^SJ46^ was produced from its native promoter on a low-copy number plasmid in *E. coli* MG1655, it decreased the efficiency of plaquing (EOP) of Bas11 by over 10^4^-fold (Fig. 1a and S1a), indicating it provided strong defense against Bas11. To identify the phage-encoded activator(s) in Bas11, we isolated spontaneous Bas11 mutants that largely overcome CapRel^SJ46^ defense (Fig. 1b and S1b). Surprisingly, although the MCP of Bas11 is highly similar (85% identical) to that of SECΦ27 (Fig. S1c), none of the Bas11 escape mutants mapped to its major capsid protein. Instead, all six escaping phage clones contained mutations in the genomic region of gene *54*, which encodes a small hypothetical protein of 66 amino acids, Gp54^Bas11^ (Fig. 1b-c). Escape clones 1 and 2 each had a single nucleotide substitution that led to a premature stop codon (W43*) or a single amino acid substitution (G24D) in Gp54^Bas11^. Clones 3 and 4 each contained a mutation immediately upstream of the gene *54* coding region, likely within its promoter, that could alter Gp54^Bas11^ levels (Fig. 1c). Lastly, clones 5 and 6 each had a large deletion encompassing gene *54* and nearby genes (Fig. 1c). These results suggested that loss-of-function mutations in gene *54* allow Bas11 to largely overcome CapRel^SJ46^ defense. Notably, Gp54^Bas11^ does not show any sequence similarity to the MCP of SECΦ27.

**Figure 1.**
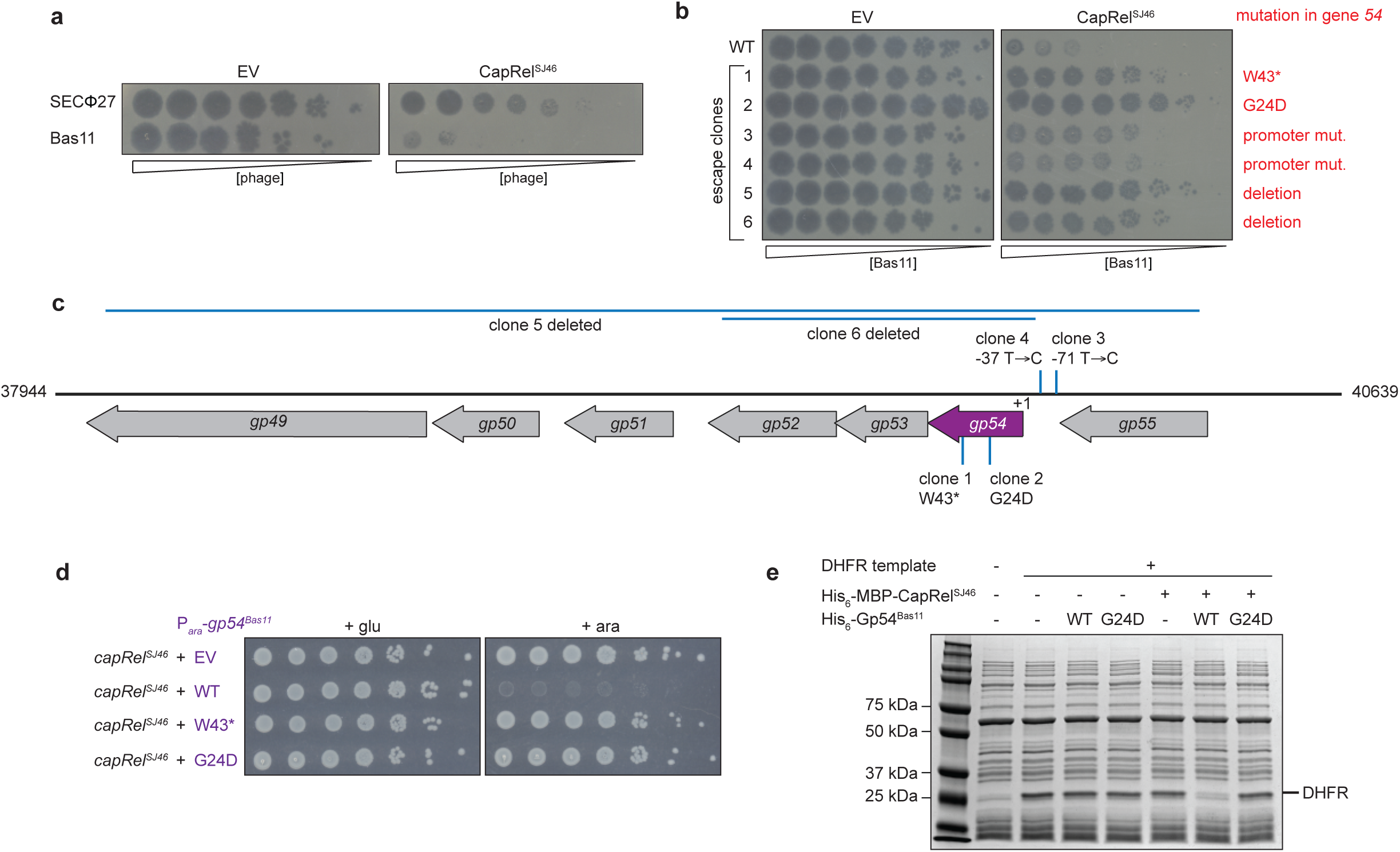
Gp54 in Bas11 is an activator of the CapRel^SJ46^ defense system. (**a**) Serial, tenfold dilutions of the indicated phages spotted on lawns of cells harboring an empty vector (EV) or a plasmid expressing CapRel^SJ46^. Relative phage concentration is indicated by the height of the wedge. (**b**) Serial dilutions of six escape clones of Bas11 and a control wild-type phage spotted on lawns of cells harboring an empty vector (EV) or a CapRel^SJ46^ expression vector, with the corresponding mutations in gene *54* labeled in red. (**c**) Schematic of the gene *54* genomic region in Bas11, with the mutations in the escape clones from (**b**) labeled. (**d**) Cell viability assessed by serial dilutions of cells expressing CapRel^SJ46^ from its native promoter and the indicated variant of Gp54^Bas11^ from an arabinose-inducible promoter on media containing glucose or arabinose. (**e**) *In vitro* transcription-translation assays using DHFR production from a DNA template as the readout. Purified His_6_-MBP-CapRel^SJ46^ and the wild-type or the G24D variant of His_6_-Gp54^Bas11^ were added to the reactions.

We hypothesized that the wild-type phage protein Gp54^Bas11^ may be an activator of CapRel^SJ46^, with the escape mutants enabling the phage to overcome defense by preventing activation. To test whether Gp54^Bas11^ is sufficient to activate CapRel^SJ46^, which blocks cell growth when active^11^, we co-produced wild-type or a mutant variant of Gp54^Bas11^ with CapRel^SJ46^ in the absence of phage infection. Wild-type Gp54^Bas11^ rendered CapRel^SJ46^ toxic, whereas neither variant (W43* or G24D) had any effect on cell growth when co-produced with CapRel^SJ46^ (Fig. 1d). As a control, we verified that wild-type and the mutant variants of Gp54^Bas11^ were not toxic on their own in the absence of CapRel^SJ46^ (Fig. S1d).

We then tested whether Gp54^Bas11^ can activate CapRel^SJ46^ to inhibit protein translation in a reconstituted *in vitro* transcription-translation system. Incubating purified His_6_-MBP-CapRel^SJ46^ with purified His_6_-Gp54^Bas11^ strongly inhibited the synthesis of a model protein, DHFR, whereas the G24D variant of Gp54^Bas11^ had no effect (Fig. 1e). We verified that the G24D variant was still properly folded as it had a circular dichroism spectrum comparable to the wild-type protein (Fig. S1e). Taken together, our results indicated that wild-type Gp54^Bas11^, like the previously identified major capsid protein from phage SECΦ27, activates CapRel^SJ46^.

### CapRel^SJ46^ directly senses Gp54^Bas11^ via its antitoxin domain

CapRel^SJ46^ consists of a conserved N-terminal toxin domain that can pyrophosphorylate tRNAs and a C-terminal antitoxin domain containing a zinc-finger-like domain (pseudo-ZFD) flanked by α-helices referred to as anchors (Fig. 2a). The antitoxin domain is highly variable among CapRel homologs (Fig. S2a) and largely determines the phage specificity of CapRel defense^11^. The antitoxin of CapRel^SJ46^ directly binds the MCP of SECΦ27, serving as a phage infection sensor^11^. To test whether Gp54^Bas11^ also interacts with CapRel^SJ46^ to activate it, we immunoprecipitated CapRel^SJ46^-FLAG from cells co-producing wild-type Gp54^Bas11^-HA or the G24D variant, after verifying that tags did not affect protein functions (Fig. S2b). We found that wild-type but not the G24D variant of Gp54^Bas11^ co-immunoprecipitated with CapRel^SJ46^ (Fig. 2b). In addition, isothermal titration calorimetry (ITC) indicated that purified Gp54^Bas11^ directly binds CapRel^SJ46^ with a K*_d_* of 800 nM in a 1:1 ratio, comparable to that previously measured for MCP^SECΦ27^ (350 nM)^11^. The interaction is entropically driven suggesting the bound state is somewhat dynamic or some region of the complex becomes disordered upon binding. The binding affinity decreased at least 20-fold for the G24D variant of Gp54^Bas11^ (Fig. 2c).

**Figure 2.**
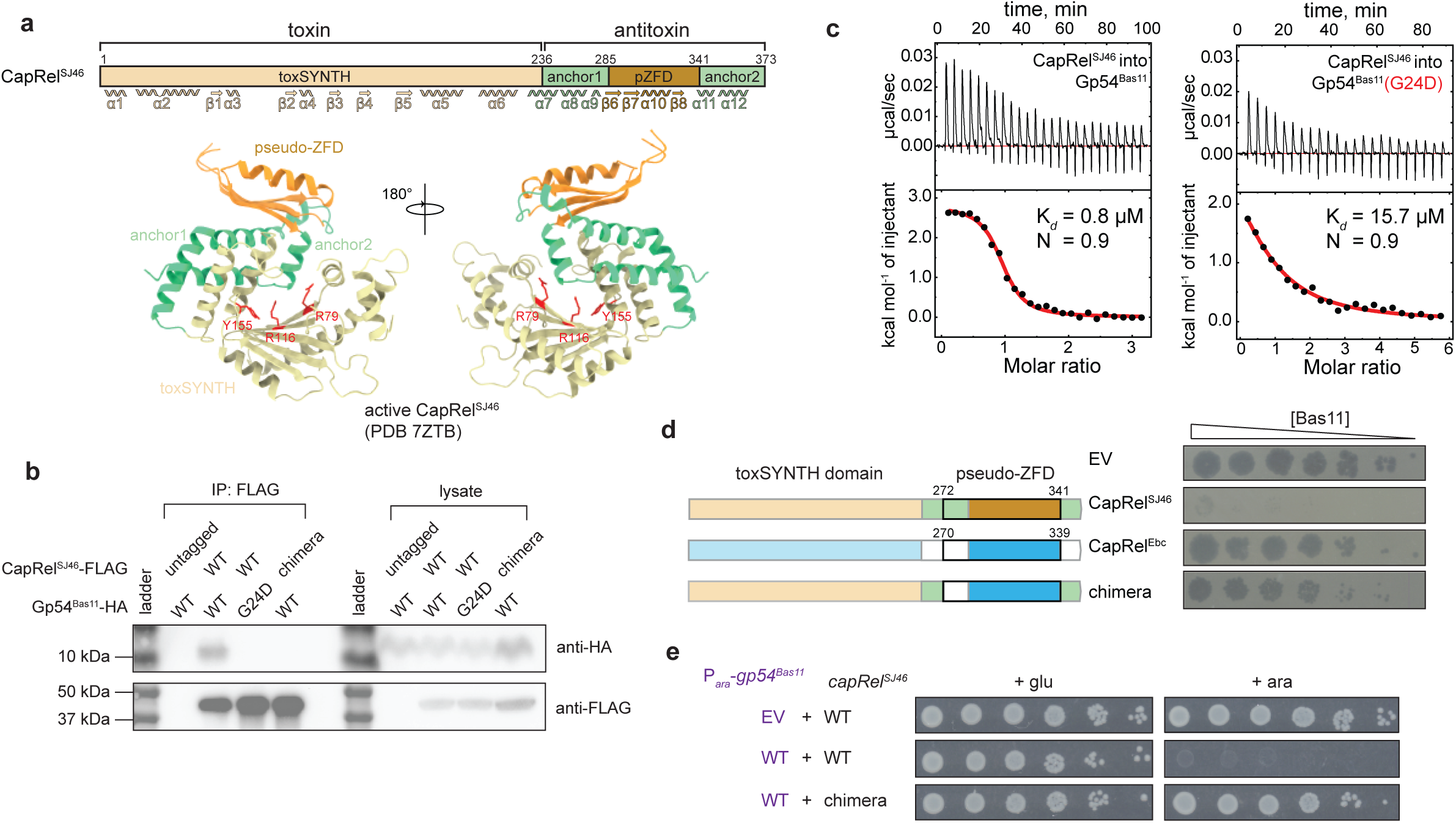
Gp54^Bas11^ binds directly to the antitoxin region of CapRel^SJ46^. (**a**) Schematic of the domain organization of CapRel^SJ46^ (*top*) and cartoon representation of the crystal structure of CapRel^SJ46^ colored by domains (*bottom*). Active site G-loop Y155 and the ATP-coordination residues R79 and R116 of the toxin domain are highlighted in red. (**b**) CapRel^SJ46^-FLAG or chimera-FLAG was immunoprecipitated from cells producing CapRel^SJ46^-FLAG or chimera-FLAG and Gp54^Bas11^-HA (wild-type or the G24D variant) and probed for the presence of the indicated Gp54^Bas11^ variant via the HA tag. Lysates used as input for the immunoprecipitation (IP) were probed as controls for expression levels. (**c**) Binding of CapRel^SJ46^ to the wild-type or G24D variant of Gp54^Bas11^ monitored by isothermal titration calorimetry (ITC) with binding affinity (K*_d_*) and stoichiometry (N) noted. (**d**) Serial dilutions of the Bas11 phage spotted on lawns of cells harboring the indicated CapRel constructs or an empty vector. *Left*, schematic of the CapRel constructs. (**e**) Serial dilutions of cells expressing CapRel^SJ46^ or the chimera from its native promoter and the wild-type Gp54^Bas11^ from an arabinose-inducible promoter on media containing glucose or arabinose.

To test whether Gp54^Bas11^ is also sensed by the antitoxin domain of CapRel^SJ46^, we spotted Bas11 phages onto cells producing the homolog CapRel^Ebc^ from *Enterobacter chengduensis* or a chimera that replaced most of the CapRel^SJ46^ antitoxin with the corresponding region of CapRel^Ebc^ (Fig. 2d and S2c). Unlike CapRel^SJ46^, neither CapRel^Ebc^ nor the chimera provided robust defense against Bas11 (Fig. 2d and S2c), despite their abilities to protect against another phage, T7 (Fig. S2d). In addition, the chimeric version of CapRel was no longer toxic to cells when co-produced with wild-type Gp54^Bas11^ (Fig. 2e), and Gp54^Bas11^ did not co-precipitate with the chimeric CapRel (Fig. 2b). These results indicated that the C-terminal antitoxin domain of CapRel^SJ46^ is important for sensing Gp54^Bas11^ despite Gp54’s lack of sequence or structural similarity to the MCP from SECΦ27.

To further investigate how the antitoxin of CapRel^SJ46^ senses Gp54^Bas11^, we mutagenized this domain through error-prone PCR and selected for CapRel^SJ46^ mutants that were no longer activated by Gp54^Bas11^. The single substitutions N275D (Fig. 3a), L270P, and L276P (Fig. S3a) each largely abolished the toxicity of CapRel^SJ46^ when co-produced with Gp54^Bas11^, and substantially weakened CapRel^SJ46^ defense against Bas11 (Fig. S3b). Notably, these residues all lie within α-helix 9, which is part of anchor-1 in the antitoxin, a highly variable region among CapRel homologs (Fig. 2a, S2a and S3c); thus, we hypothesized that α9, formed by residues 270-279, might be critical for CapRel^SJ46^ to interact with Gp54^Bas11^. To further probe the role of anchor-1 in interacting with Gp54^Bas11^, we made substitutions in other non-conserved residues within α9 and tested their activation by Gp54^Bas11^. The substitutions K278E (Fig. 3a), D273K and S279P (Fig. S3a) each reduced or abolished the toxicity of CapRel^SJ46^ following induction of Gp54^Bas11^, supporting a key role for this helix in sensing Gp54^Bas11^.

**Figure 3.**
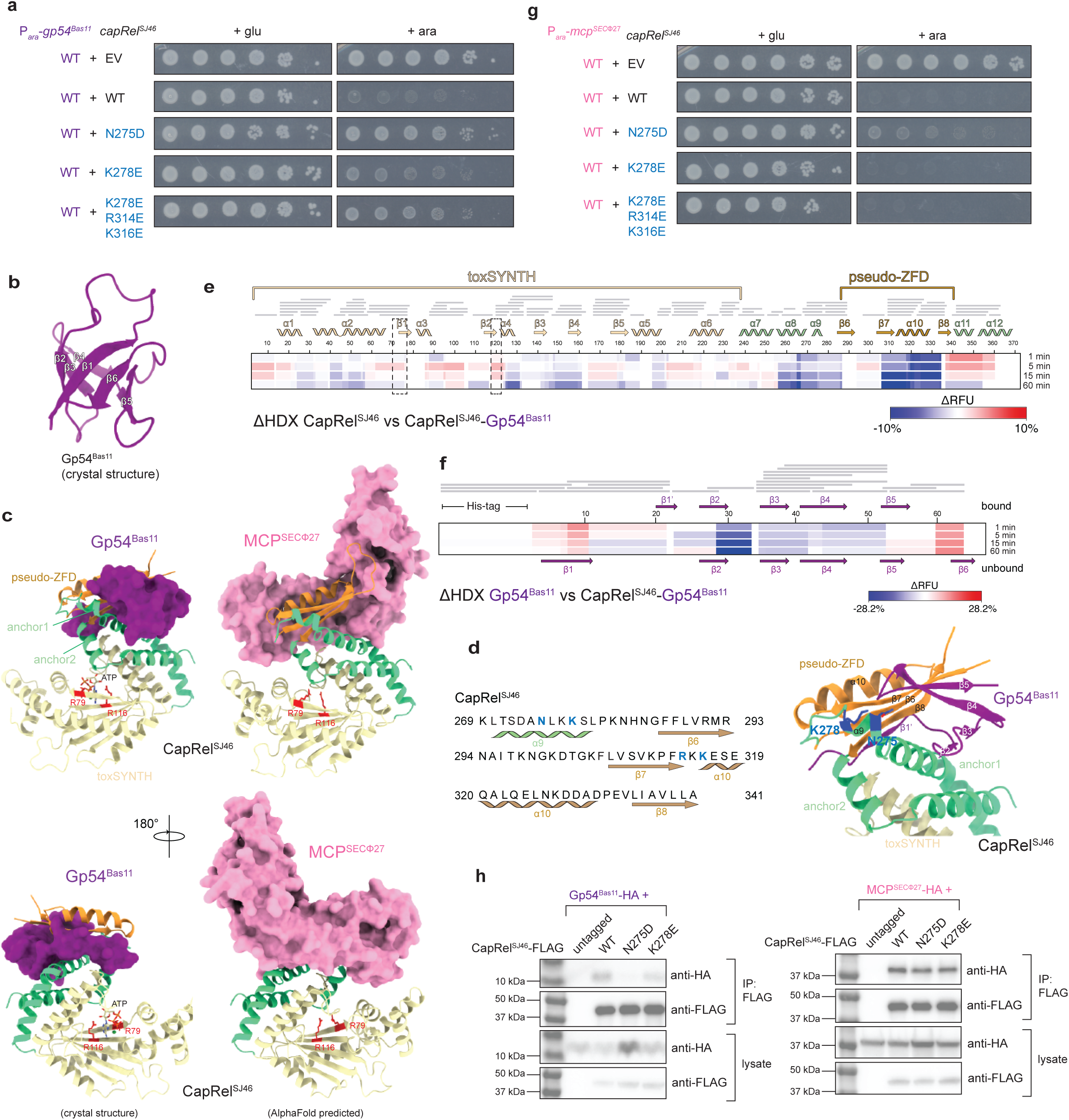
Gp54^Bas11^ and MCP^SECΦ27^ bind overlapping but distinct regions of CapRel^SJ46^. (**a**) Serial dilutions of cells expressing the indicated variant of CapRel^SJ46^ from its native promoter and the wild-type Gp54^Bas11^ from an arabinose-inducible promoter on media containing glucose or arabinose. (**b**) Cartoon representation of the crystal structure of Gp54^Bas11^. (**c**) *Left*, crystal structure of the complex of Gp54^Bas11^ (purple) bound to CapRel^SJ46^ (colored by domains). *Right*, structural model of the complex of CapRel^SJ46^ and MCP^SECΦ27^ (pink) by AlphaFold. ATP coordination residues of CapRel^SJ46^ toxin domain are highlighted in red. (**d**) *Left*, amino acid sequence of the antitoxin region of CapRel^SJ46^ with the residues substituted colored. *Right*, details of the interface formed by the antitoxin domain of CapRel^SJ46^ and Gp54^Bas11^ (purple) with the residues substituted colored in blue. (**e**) Differential HDX (ΔHDX) between CapRel^SJ46^ and CapRel^SJ46^-Gp54^Bas11^ displayed as a difference heat map. Change in relative fractional units (ΔRFU) are color coded with red indicating increased deuteration of CapRel^SJ46^ in the presence of Gp54^Bas11^ and blue indicating lower deuteration. Grey bars indicate peptides identified in mass spectrometry analysis. The region corresponding to the toxin domain (toxSYNTH) active site is highlighted by dashed-line boxes. (**f**) Same as in (**e**) but comparing Gp54^Bas11^ and CapRel^SJ46^-Gp54^Bas11^. (**g**) Same as in (**a**) but with the wild-type MCP from SECΦ27. (**h**) CapRel^SJ46^-FLAG or the indicated variant was immunoprecipitated from cells producing CapRel^SJ46^-FLAG and Gp54^Bas11^-HA or MCP^SECΦ27^-HA and probed for the presence of Gp54^Bas11^ or MCP^SECΦ27^ via the HA tag.

### Gp54^Bas11^ changes conformation and partially unfolds upon binding to CapRel^SJ46^

To gain better structural insight into the interaction between CapRel^SJ46^ and Gp54^Bas11^, we first solved a crystal structure of Gp54^Bas11^ to 2.3 Å resolution (Fig. 3b, S3d and Table S1). This structure revealed a small, 6-stranded β-barrel with one prominent loop between β-strands β1 and β2. β-barrels with this topology are very rare in nature^19^. The closest structural homologs of Gp54^Bas11^ are 5-stranded β-barrel SH3 domains (DALI Z-score of 3.8) with a 3_10_ α-helix replacing the additional β-strand (Fig. S3e). Thus, the structure of Gp54^Bas11^ is significantly distinct from that of the MCP from SECΦ27, with RMSD greater than 14 Å (Fig. S3f).

Next, we determined the structure of a CapRel^SJ46^-Gp54^Bas11^ complex to 2.2 Å resolution (Fig. 3c, S4a-c and Table S1). This complex had ATP bound in the pyrophosphate donor site (Fig. 3c and S4c), stacked between R79 and R116, similar to that of other RelA/SpoT homolog (RSH) enzymes^20,21^, which indicated that the complex captures the active state of the enzyme. The complex structure revealed that Gp54^Bas11^ interacts with the pseudo-ZFD and anchor-1 of the antitoxin, with an interface of ∼1650 Å^2^ that is partially overlapping, but largely distinct from that formed between CapRel^SJ46^ and MCP^SECΦ27^ (Fig. 3c). In this triggered and active state, residue Y355 in CapRel^SJ46^, which is part of the YXXY motif that normally blocks the ATP-binding site in the closed state^11^, is tethered to Gp54^Bas11^ β2 via K269 and cannot interact with the toxin domain (Fig. S4d). Small angle X-ray scattering (SAXS) analysis of the CapRel^SJ46^-Gp54^Bas11^ complex was compatible with the crystal structure and indicated, by comparison to the unbound CapRel^SJ46^, that Gp54^Bas11^ effectively clamps the pseudo-ZFD to both anchors, precluding recoil toward the toxin active site (Fig. S4e-f and Table S2).

Each of the substitutions in CapRel^SJ46^ identified above as affecting activation (Fig. 3a and S3a-b) map to the interface formed with Gp54^Bas11^ (Fig. 3d and S5a). In the complex, the hydrophobic core of the Gp54^Bas11^ β-barrel binds to the amphipathic anchor-1 and to the pseudo-ZFD of CapRel^SJ46^. In particular, W43 of Gp54^Bas11^ contacts β6 and β7 from the pseudo-ZFD and residues G24, I25, S39, and L41 from Gp54^Bas11^, while W53 of Gp54^Bas11^ becomes embedded between L270 and L276 of anchor-1 in CapRel^SJ46^ and I29 and L31 of Gp54^Bas11^ (Fig. S5a-b). The interface is further stabilized by a polar network between D273, N275, K278 and S279 from CapRel^SJ46^ and Q47 and N50 from Gp54^Bas11^ (Fig. S5a).

Remarkably, binding of Gp54^Bas11^ to CapRel^SJ46^ involves a significant topological rearrangement (Fig. S5c). In the complex, the β-barrel of Gp54^Bas11^ unfolds, with β1 and β6 becoming disordered, consistent with the entropy-driven binding suggested by ITC (Fig. 2c). While β2-β5 bind on one side of the pseudo-ZFD of CapRel^SJ46^ interacting with β7 and anchor-1, the long β1-β2 loop of Gp54^Bas11^ folds into β-strand β1’ and binds on the other side, making contacts with β8 of the pseudo-ZFD (Fig. 3d and S4a). These interactions produce a ’hybrid’ 8-stranded antiparallel, twisted β-sheet comprised of β-strands from both Gp54^Bas11^ and CapRel^SJ46^ that wraps around anchor-1 of CapRel^SJ46^ (Fig. 3d and S4a). Additionally, β1’ of Gp54^Bas11^ binding to CapRel^SJ46^ β8, moves the pseudo-ZFD further from the active site compared to the unbound open state of CapRel^SJ46^, which likely primes the enzyme to bind and modify target tRNAs (Fig. S5d). Finally, we noted that the N-terminal region of Gp54^Bas11^ β1’ (and possibly the disordered N-terminus) also makes contact with the cap of α10 (residues 314-317) in CapRel^SJ46^ (Fig. S5e). Supporting the relevance of this interaction, we found that the substitutions R314E and K316E, together with K278E, further reduced the toxicity of CapRel^SJ46^ following induction of Gp54^Bas11^ (Fig. 3a).

To further validate the ordered-to-disordered transition of Gp54^Bas11^ and binding interface of the CapRel^SJ46^-Gp54^Bas11^ complex, we used hydrogen–deuterium exchange (HDX) coupled to mass spectrometry to compare the CapRel^SJ46^-Gp54^Bas11^ complex with the unbound proteins. This analysis revealed protection of anchor-1 and the pseudo-ZFD regions of CapRel^SJ46^, particularly of β7-β8 and α8-α10 (Fig. 3e and S6a-d), which almost perfectly matched the crystallographic interface. We also observed an increase in deuterium uptake by anchor-2 (α11 and α12), which contains the YXXY motif that interacts with the toxin active site (Fig. 3e and S6d). This deprotection reflects the opening of CapRel^SJ46^ upon binding Gp54^Bas11^. On the Gp54^Bas11^ side, the newly formed β1’ and the β2-β4 region showed decreases in deuterium uptake, again consistent with the crystal structure (Fig. 3f). The HDX-MS also showed an increase in uptake in the N- and C-terminal regions of Gp54^Bas11^ (β1 and β6 in the unbound state), consistent with unfolding of the β-barrel upon binding CapRel^SJ46^ and disorder in these regions in the crystal structure. Collectively, our results demonstrate that a dynamic, entropically favorable bound state drives the activation of CapRel^SJ46^ by Gp54^Bas11^.

### Activation of CapRel^SJ46^ by Gp54^Bas11^ and the MCP^SECΦ27^ can be genetically separated

Importantly, the α-helix (a9) in anchor-1 of CapRel^SJ46^ that makes extensive contacts with Gp54^Bas11^ does not contribute significantly to the interface with MCP^SECΦ27^ (Fig. 3c). Thus, we hypothesized that the substitutions in this region of CapRel^SJ46^ that disrupt activation by Gp54^Bas11^ would not impact activation by MCP^SECΦ27^. To test this hypothesis, we co-produced our CapRel^SJ46^ variants with MCP^SECΦ27^ and found that the single substitutions N275D and K278E, as well as the triple substitution K278E R314E K316E, did not substantially affect activation by MCP^SECΦ27^ despite their reduced activation by Gp54^Bas11^ (Fig. 3g). We also found that the N275D and K278E variants of CapRel^SJ46^ co-precipitated with MCP^SECΦ27^ as well as the wild-type CapRel^SJ46^, but, as expected based on the crystal structure (Fig. 3c-d), had reduced binding to Gp54^Bas11^ in this assay (Fig. 3h).

Taken all together, our results indicated that the antitoxin domain of CapRel^SJ46^ is critical for sensing both the MCP of SECΦ27 and Gp54 from Bas11, with overlapping but not identical regions of the antitoxin involved in the two different interactions. More broadly, these findings reveal the remarkable versatility of a zinc-finger-like domain in recognizing different proteins, which enables a single bacterial defense protein to sense multiple phage-encoded activators.

### Gp54^Bas11^ homologs in other related phages do not activate CapRel^SJ46^

Given that both the MCP of SECΦ27 and Gp54 from Bas11 can activate CapRel^SJ46^, we decided to examine a set of phages from the BASEL collection – Bas5, Bas8, and Bas10 – that are closely related to SECΦ27 and Bas11 and that encode homologs of the MCP (Fig. S1c) and of Gp54 (Fig. 4a). The region of the MCP shown in SECΦ27 to mediate an interaction with CapRel^SJ46^ was nearly identical in each of these phages (Fig. S1c). When examining the Gp54 homologs, we noted that those from Bas11 and Bas10 were nearly identical, with only three amino acid differences, whereas the SECΦ27, Bas5 and Bas8 homologs contained more substitutions relative to the Bas11 homolog (Fig. 4a).

**Figure 4.**
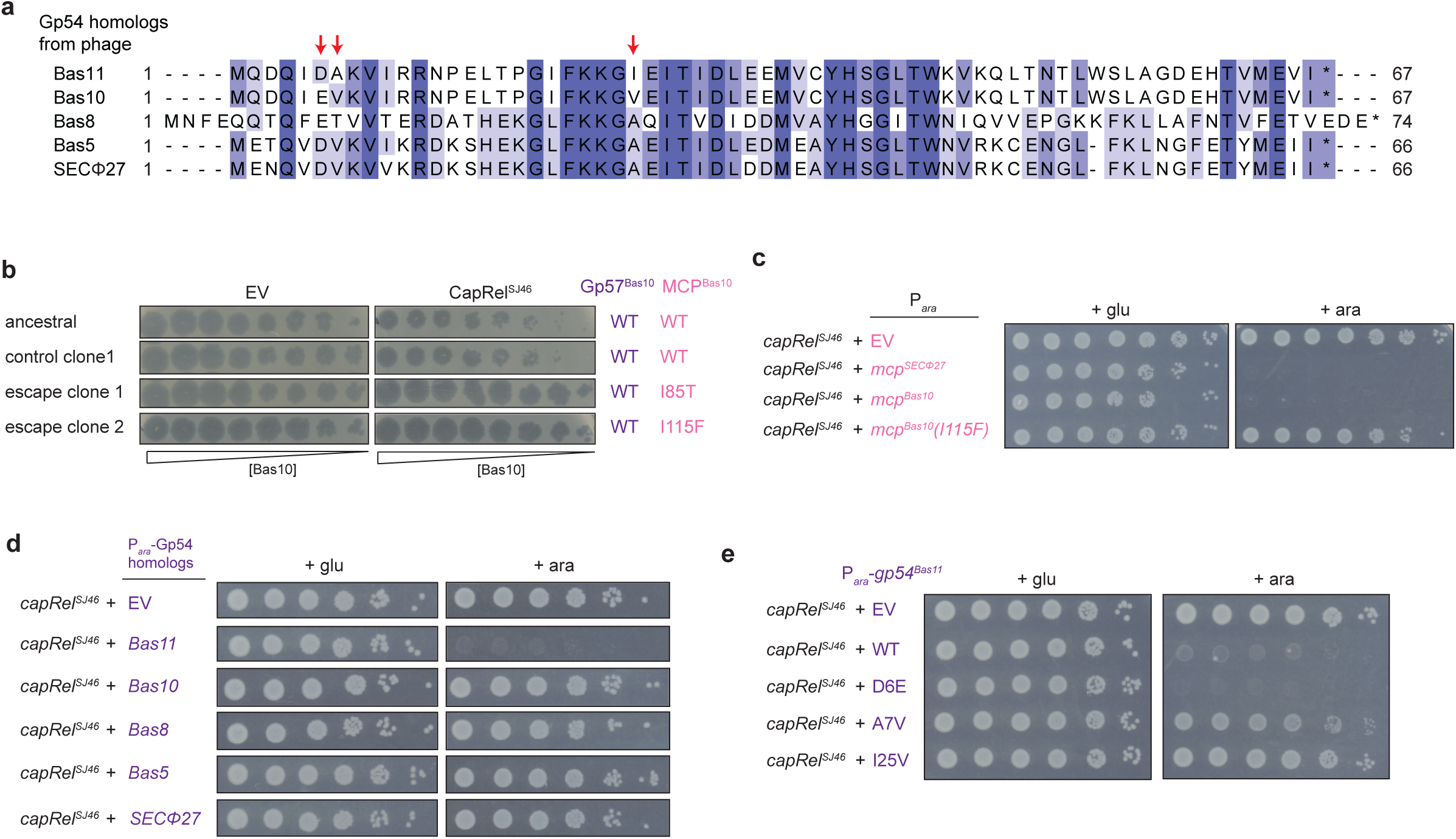
Homologs of Gp54^Bas11^ do not trigger CapRel^SJ46^. (**a**) Multiple sequence alignment of Gp54 homologs from the indicated phages. Residues that are different between the Gp54 homolog from phage Bas11 and Bas10 are labeled by red arrows. (**b**) Serial dilutions of wild-type (ancestral or from a control population evolved without selective pressure to evade CapRel^SJ46^) and the escaping clones of Bas10 spotted on lawns of cells harboring an empty vector (EV) or a plasmid producing CapRel^SJ46^. The corresponding genotype of its MCP is indicated on the right. (**c**) Serial dilutions of cells expressing CapRel^SJ46^ from its native promoter and the indicated MCP or its variant from an arabinose-inducible promoter on media containing glucose or arabinose. (**d**) Same as in (**c**) but with each Gp54 homolog from the indicated phages. (**e**) Serial dilutions of cells expressing CapRel^SJ46^ from its native promoter and the indicated variant of Gp54^Bas11^ from an arabinose-inducible promoter on media containing glucose or arabinose.

We previously showed that CapRel^SJ46^ can defend against Bas5 and Bas8, like SECΦ27, by sensing their major capsid proteins^11^. The MCPs of Bas5 and Bas8 are sufficient to activate CapRel^SJ46^ on their own, and mutations in the MCPs allowed the phages to escape defense^11^. Here, we found that CapRel^SJ46^ also defends against Bas10, reducing plaquing ∼10-fold, and selection for complete escape led to the identification of clones producing a single amino-acid substitution (I85T or I115F) in its major capsid protein, MCP^Bas10^ (Fig. 4b and S7a). Additionally, we found that wild-type MCP from Bas10, but not the I115F variant, caused toxicity to cells co-producing CapRel^SJ46^ (Fig. 4c). Notably, the same I115F substitution emerged when phage SECΦ27 was evolved to overcome CapRel^SJ46^ defense^11^.

Our results indicated that the MCP of Bas10, like that of Bas5, Bas8, and SECΦ27, is necessary and sufficient to activate CapRel^SJ46^. However, as noted, these four phages also harbor homologs of the unrelated, alternative activator Gp54 from Bas11. We therefore tested whether the Gp54 homologs of these phages can also activate CapRel^SJ46^. However, none of the Gp54 homologs from SECΦ27, Bas5, Bas8, and Bas10 caused cellular toxicity when co-produced with CapRel^SJ46^ (Fig. 4d). These findings were consistent with our results demonstrating that mutations in the MCP-encoding gene of these phages enabled complete escape from CapRel^SJ46^ defense. We concluded that phages SECΦ27, Bas5, Bas8, and Bas10 contain only a single protein activator of CapRel^SJ46^ (their major capsid proteins), despite encoding homologs of the Gp54 activator found in Bas11 phage.

This conclusion was most surprising for Bas10, which is the most closely related to Bas11 and encodes a homolog of Gp54^Bas11^ containing only 3 amino acid differences (Fig. 4a). To test the importance of these three residues for activation of CapRel^SJ46^ by Gp54^Bas11^, we made three single substitutions (D6E, A7V, I25V) in Gp54^Bas11^ to individually introduce the residues found at these positions in the Bas10 homolog. The substitutions A7V and I25V in Gp54^Bas11^ each largely abolished its ability to activate CapRel^SJ46^ when co-produced, whereas D6E had no effect (Fig. 4e). We also made the reciprocal, individual substitutions (V7A or V25I) in the Bas10 homolog of Gp54 but found that neither substitution alone enabled activation of CapRel^SJ46^ whereas the double mutant was sufficient to activate (Fig. S7b). I25 is found in the CapRel^SJ46^ - Gp54^Bas11^ complex interface, as is the adjacent G24 (Fig. S7c and S5b). The substitutions I25V and G24D (identified as an escape mutant) likely disrupt binding and thereby abolish activation. In contrast, A7 is disordered in the complex but part of the β-barrel core in unbound Gp54^Bas11^ (Fig. S7d). The A7V substitution may have stabilized the unbound β-barrel, which would also prevent activation by preventing the unfolding of Gp54^Bas11^.

### Bas11 encodes two activators of CapRel^SJ46^

The results presented thus far raised a conundrum: our escape mutant screen with Bas11 revealed only mutations in gene *54* (Fig. 1b) and Gp54^Bas11^ was sufficient to activate CapRel^SJ46^ (Fig. 1d), but the major capsid protein of Bas11 is identical to that of Bas10 where the MCP is the sole trigger for CapRel^SJ46^. We revisited our Bas11 escape mutants and noted that each only partially escaped CapRel^SJ46^ defense with about 10-100 fold reduction in EOP (Fig. 1b and S1b). Even the clone of Bas11 with a deletion of gene *54* (clone 5 in Fig. 1b) still formed smaller plaques when spotted onto cells containing CapRel^SJ46^ compared to cells with an empty vector (Fig. 1b). We tried to evolve this mutant clone of Bas11 to completely overcome CapRel^SJ46^ defense, and succeeded in isolating mutants that now plaqued the same on CapRel^SJ46^-containing cells as empty vector cells (Fig. 5a and S8a). Strikingly, whole phage genome sequencing revealed that all of our escape mutants produced an I115F substitution in the major capsid protein, MCP^Bas11^, in addition to the deletion of the gene *54* region (Fig. 5a). As shown above, the wild-type MCP from Bas11 (which is identical to that of Bas10) was sufficient to activate CapRel^SJ46^ and the substitution I115F completely ablated this activation (Fig. 4c). We engineered wild-type Bas11 phage to encode only the I115F substitution in its MCP, *i.e.* with gene *54* present, and observed that this substitution alone was also insufficient for Bas11 to completely escape CapRel^SJ46^ defense with a 10-fold reduction in EOP and smaller plaques (Fig. 5b and S8b). Thus, our results demonstrate that Bas11 encodes two activators of CapRel^SJ46^, and it can only fully escape CapRel^SJ46^ defense when both activators are mutated.

**Figure 5.**
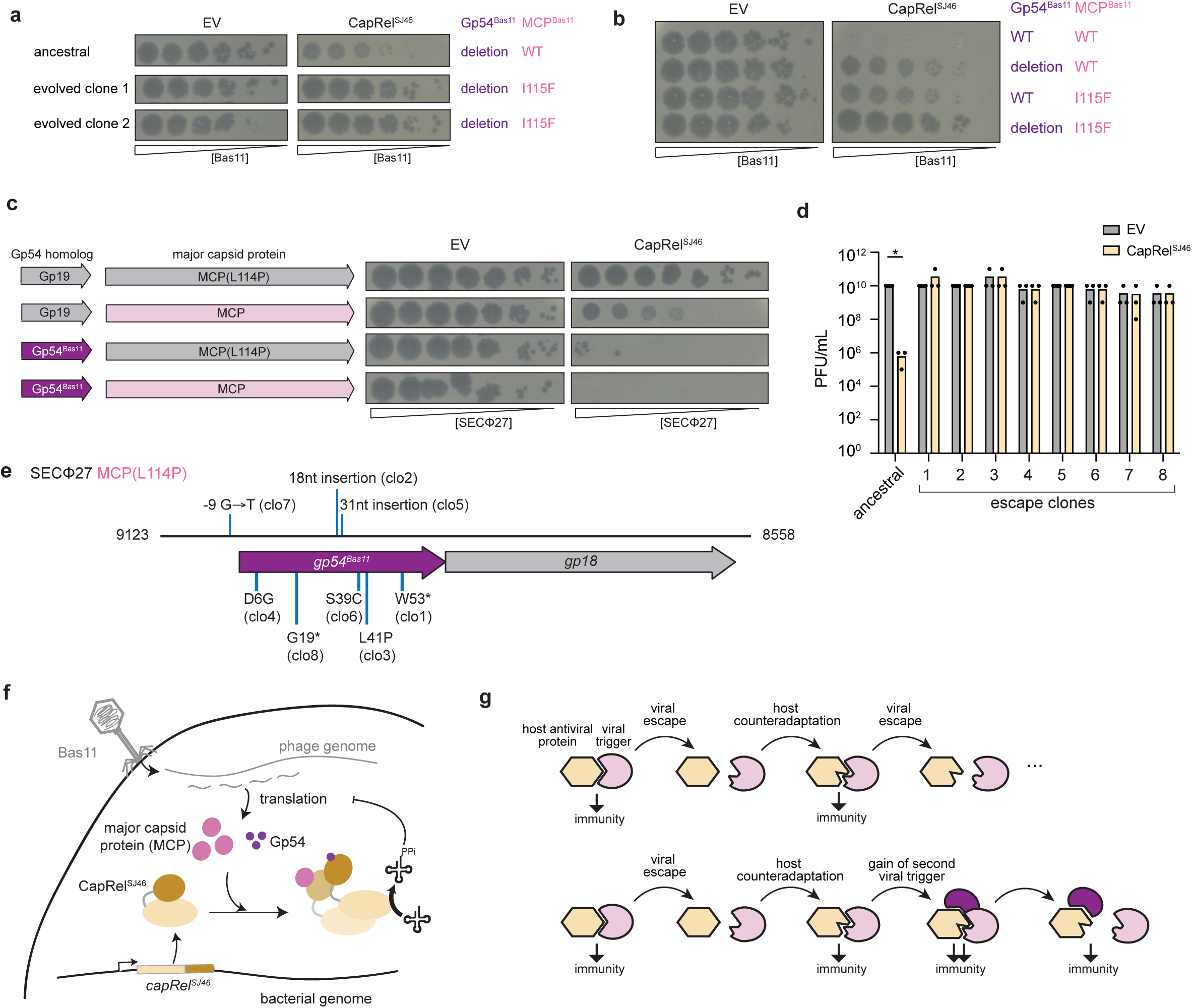
CapRel^SJ46^ can sense and respond to two unrelated trigger proteins in Bas11 phage. (**a**) Serial dilutions of the ancestral and evolved Bas11 phage spotted on lawns of cells harboring an empty vector (EV) or a plasmid expressing CapRel^SJ46^. The corresponding genotypes of its MCP or Gp54 were indicated on the right. (**b**) Serial dilutions of the indicated Bas11 phage spotted on lawns of cells harboring an empty vector (EV) or a plasmid expressing CapRel^SJ46^. The corresponding genotypes of its MCP or Gp54 were indicated on the right. (**c**) *Left*, schematics of the region encoding Gp19 (the Gp54 homolog from SECΦ27) or the MCP. *Right*, serial dilutions of the indicated SECΦ27 phage spotted on lawns of cells harboring an empty vector (EV) or a plasmid expressing CapRel^SJ46^. (**d**) Summary of plaque forming units (PFU) obtained for the ancestral SECΦ27 MCP(L114P) phage carrying Gp54^Bas11^ or eight escape clones after spotting onto cells producing CapRel^SJ46^ or harboring an empty vector (EV). 3 independent replicates are shown. Asterisks indicate p < 0.05 (unpaired two-tailed t-test). (**e**) Schematic of the gene *19* genomic region in SECΦ27 MCP(L114P) phage replaced by gene *54* from Bas11, with the mutations in the escape clones from (**d**) labeled. (**f**) Model for the activation of CapRel^SJ46^ by the major capsid protein (MCP) and Gp54 in phage Bas11. (**g**) *Top*, schematic of the Red Queen dynamic between an anti-viral immunity protein such as CapRel^SJ46^ and its viral trigger proteins, following the schematic in ref^25^. When a single protein is recognized, successive rounds of viral escape and host counteradaptation produce a conventional coevolutionary dynamic. *Bottom*, when two proteins are recognized by an antiviral system like CapRel^SJ46^, viral escape is more difficult, as any single escape mutation will not prevent activation of immunity.

To further compare the activation of CapRel^SJ46^ by the MCP and Gp54^Bas11^, we engineered phage SECΦ27 such that it encodes one or both proteins as activators. As shown previously, despite encoding a Gp54^Bas11^ homolog (called Gp19), wild-type SECΦ27 normally triggers CapRel^SJ46^ only via its MCP, with a single substitution in the MCP (L114P) allowing the phage to completely escape defense. We replaced the coding sequence of the SECΦ27 homolog (Gp19) with that of Gp54^Bas11^ in both wild-type SECΦ27 and the SECΦ27 MCP(L114P) escape phage (Fig. 5c). When Gp54^Bas11^ was introduced into SECΦ27 MCP(L114P), it restored CapRel^SJ46^-dependent defense, with a > 10^5^-fold decrease in EOP (Fig. 5c and S8c). Defense against this phage, which uses Gp54^Bas11^ as the activator of CapRel^SJ46^, was stronger than defense against wild-type SECΦ27 which produces ∼10^2^-fold reduction in EOP and only uses the MCP to trigger CapRel^SJ46^ (Fig. 5c and S8c). However, we found that some clones of this engineered phage spontaneously escaped CapRel^SJ46^ defense (Fig. 5c). We isolated eight such clones that completely overcame defense and found that each harbored a mutation in the region encoding its activator Gp54^Bas11^ (Fig. 5d-e and S8d). Three clones had a single amino-acid substitution (D6G, S39C, or L41P) in Gp54^Bas11^, and these variants no longer activate CapRel^SJ46^ (Fig. S8e). Notably, S39 and L41 are part of the hydrophobic pocket involved in interacting with CapRel^SJ46^ (Fig. S8f and S5b) in the crystal structure. D6 is disordered in the complex, so the D6G substitution may disrupt an interaction not captured in the crystal structure or stabilize the unbound state of Gp54^Bas11^ to prevent CapRel^SJ46^ activation.

Finally, when Gp54^Bas11^ was introduced to wild-type SECΦ27 such that both wild-type MCP and Gp54^Bas11^ were present in the SECΦ27 genome, it led to a stronger defense phenotype (> 10^6^-fold reduction in EOP) compared to phages encoding a single activator, and no spontaneous escape mutants were observed (Fig. 5c and S8c). These results indicated that Gp54^Bas11^ functioned as a potent activator when introduced into a related phage SECΦ27, and harboring both activators (Gp54^Bas11^ and the MCP) in its genome rendered this phage extremely sensitive to CapRel^SJ46^ defense, as with the native Bas11 phage. By sensing two activators encoded in a single phage genome, CapRel^SJ46^ can provide strong defense and limit the ability of phages to escape defense.

## Discussion

Previously, we demonstrated that CapRel^SJ46^, a fused toxin-antitoxin system, provides *E. coli* with robust anti-phage defense by sensing the major capsid proteins of many phages^11^ (Fig. 5f). Upon infection by phages SECΦ27, Bas5, Bas8 and Bas10, their newly synthesized MCPs directly bind to the C-terminal antitoxin domain of CapRel^SJ46^ to stabilize an active, open conformation. Activated CapRel^SJ46^ then pyrophosphorylates tRNAs to inhibit translation and restrict viral propagation through the bacterial population. Although sensing an essential and conserved component of the phage, such as the MCPs, limits the ability of phages to overcome a defense system, phages can still escape defense by mutating its trigger protein. Thus, it may be advantageous for the defense system to sense more than one component of the phage.

Here, we discovered an additional protein trigger for CapRel^SJ46^ in the phage Bas11, besides the MCP. This alternative trigger, Gp54^Bas11^, is a small protein of 66 amino acids with unknown function. Despite lacking sequence and structural similarity to the MCPs, Gp54^Bas11^ can bind to the C-terminal antitoxin domain of CapRel^SJ46^ and directly activate it (Fig. 5f). Combining structural studies, mutagenesis, and biochemical analyses, our work indicates that Gp54^Bas11^ directly binds the antitoxin domain of CapRel^SJ46^, like the MCPs. The interfaces used by the two protein activators (MCP and Gp54^Bas11^) overlap but also involve distinct regions. These findings highlight the versatility of a zinc-finger-like domain in interacting with multiple, structurally different proteins. Using such a promiscuous, yet still selective domain as a phage-infection sensor enables a single bacterial defense protein to respond to more than one phage trigger.

Unlike the major capsid protein, which is a conserved and essential structural element of the phage, Gp54^Bas11^ is a small protein of unknown function that is not essential to phage Bas11 under laboratory conditions. Gp54^Bas11^ might benefit the phage in the wild, possibly by inhibiting another defense system. Recent work has identified other small, non-essential phage proteins that activate one anti-phage defense system, while also inhibiting another defense system. For example, the Ocr protein of phage T7 inhibits restriction-modification systems, but can also activate the PARIS defense system^22–24^.

Analogous to PAMP-based sensing in eukaryotic innate immunity, recent work has identified phage-encoded triggers of several anti-phage defense systems in bacteria^7–11,17^. Different systems have been shown to recognize diverse phage proteins or nucleic acids. However, only a single direct trigger has been identified for a given defense system and it is often implied that defense systems and triggers have one-to-one relationships. Our work now demonstrates that CapRel^SJ46^ has evolved to directly and simultaneously, *i.e.* during a single infection, detect two different proteins (MCP and Gp54^Bas11^) from phage Bas11 (Fig. 5f). The detection of multiple phage factors is likely a common feature of bacterial immunity proteins. For instance, a high-throughput screen for bacteria-encoded triggers of an anti-phage retron system identified multiple genes from prophages^17^. Each was sufficient, when overproduced, to trigger the retron system, but whether they each contribute to activation during infection remains to be shown. Similarly, multiple phage proteins other than Ocr can stimulate the PARIS defense system, though whether they bind directly and whether the binding is similar to or different than Ocr is not yet known^24^.

The ability to sense multiple proteins may offer at least three advantages to bacterial immune systems. One, for phages that produce multiple triggers, the immunity system can provide stronger defense than if it sensed only one protein. Two, dual sensing makes it more difficult for the phage to completely evade defense completely unless both triggers are mutated. Three, sensing multiple phage proteins may enable protection against a broader set of phages, some of which encode only one trigger or the other. Given these advantages, we anticipate that many anti-phage defense systems may have evolved similar versatility and can also directly sense multiple proteins. More broadly, sensing multiple phage proteins likely leads to complex coevolutionary dynamics between bacteria and their phage predators (Fig. 5g). The Red Queen dynamic underpinning host-pathogen relationships is often portrayed as the successive coevolution of two interacting proteins^25^, but may, in fact, involve multiple proteins stemming from many-to-one relationships between triggers and immunity proteins.

## Supplementary Figure Legends

**Figure S1.**
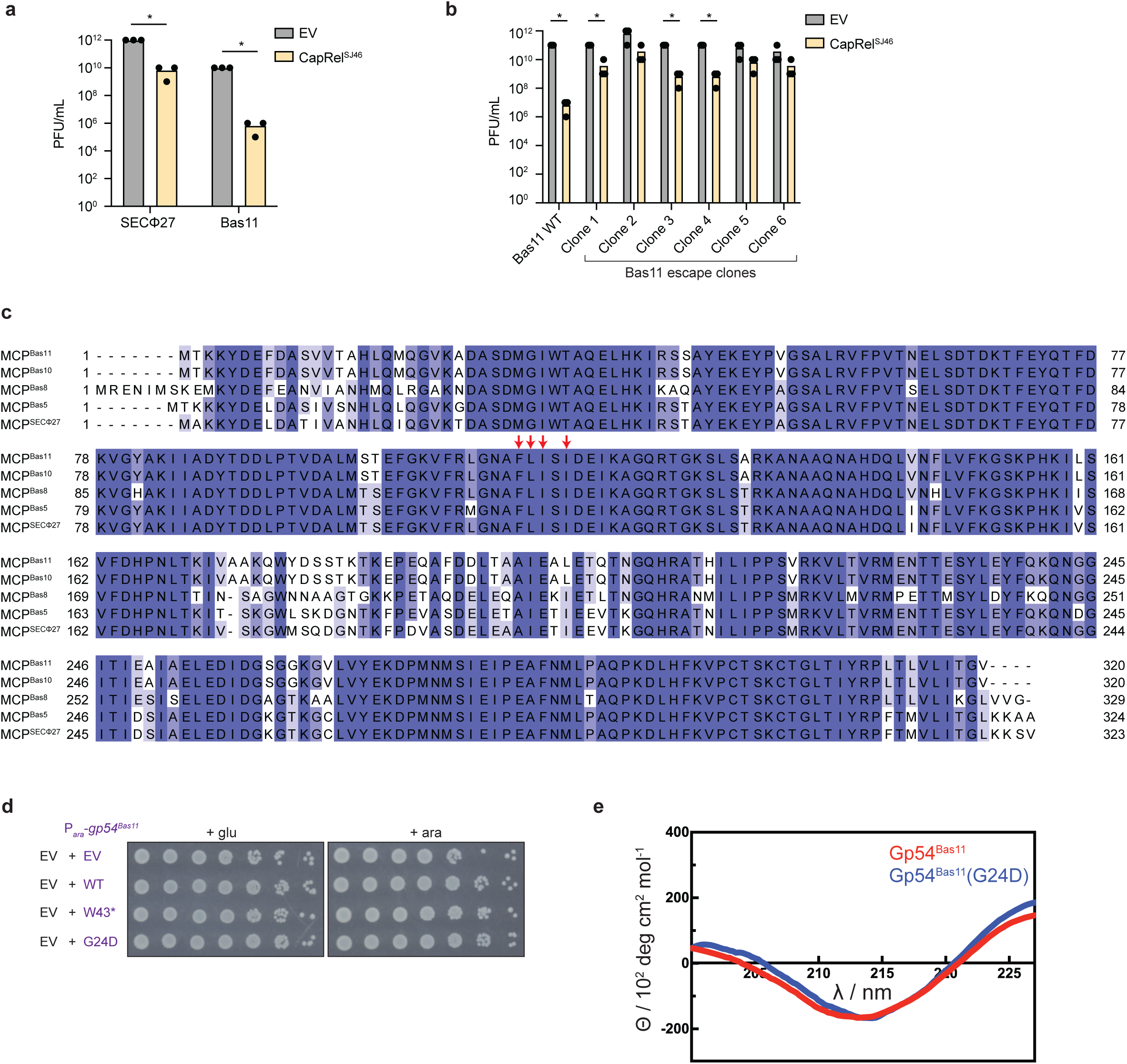
Gp54^Bas11^ activates CapRel^SJ46^. (**a**) Summary of 3 independent replicates of phage spotting assay in Fig. 1a. Asterisks indicate p < 0.05 (unpaired two-tailed t-test). (**b**) Summary of 3 independent replicates of phage spotting assay in Fig. 1b. Asterisks indicate p < 0.05 (unpaired two-tailed t-test). (**c**) Multiple sequence alignment of the major capsid proteins from phages SECΦ27, Bas5, Bas8, Bas10 and Bas11. Residues shown to be important for interaction with CapRel^SJ46^ are labeled by red arrows. (**d**) Cell viability assessed by serial dilutions of cells expressing an empty vector (EV) and the indicated variant of Gp54^Bas11^ from an arabinose-inducible promoter on media containing glucose or arabinose. (**e**) Circular dichroism spectra for purified His_6_-Gp54^Bas11^ and His_6_-Gp54^Bas11^(G24D).

**Figure S2.**
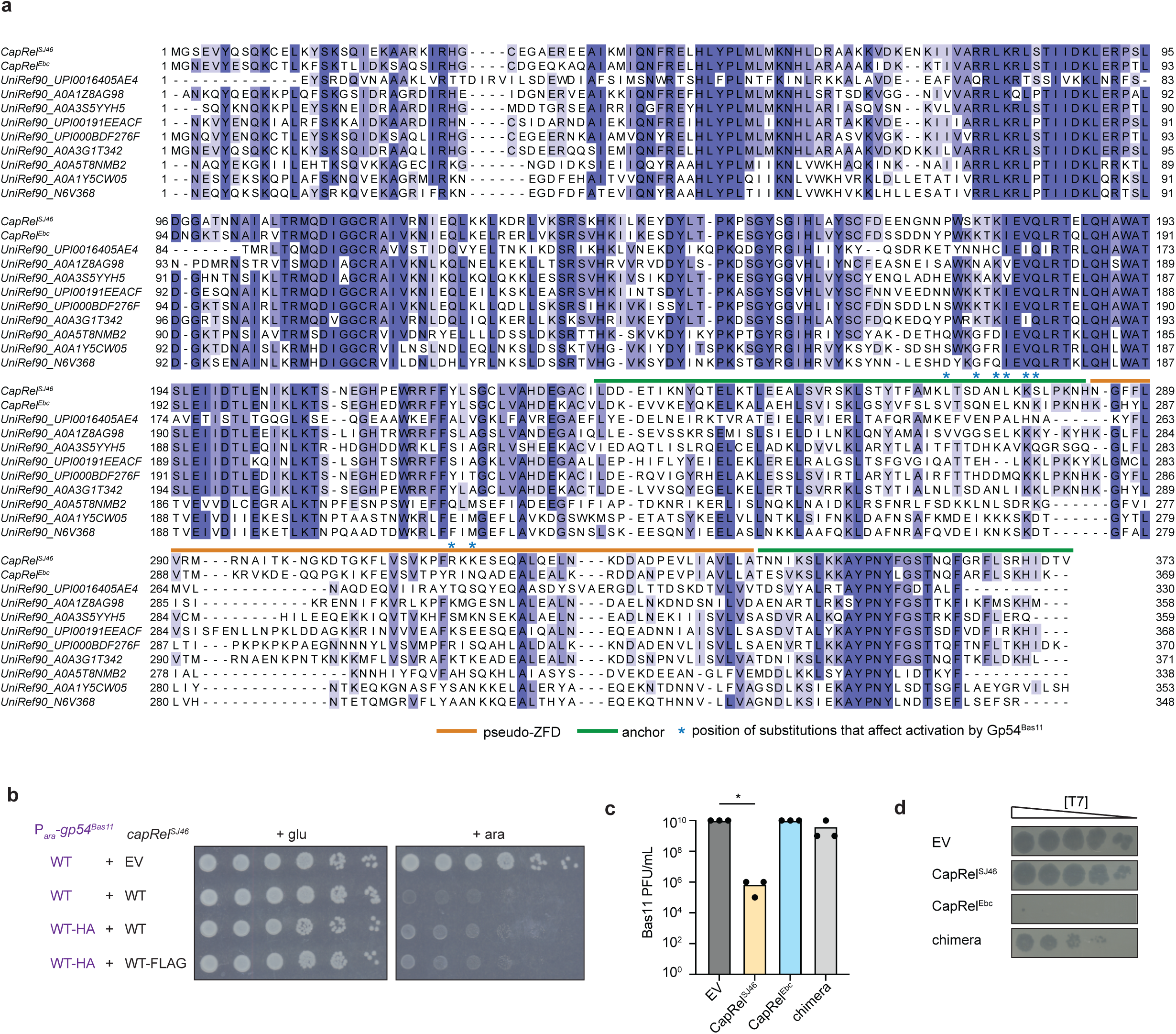
Gp54^Bas11^ binds directly to CapRel^SJ46^. (**a**) Multiple sequence alignment of CapRel^SJ46^ and diverse CapRel homologs, with labels indicating the antitoxin domain (pseudo-ZFD and anchors). The residues whose substitutions affect the activation of CapRel^SJ46^ by Gp54^Bas11^ are highlighted with stars. Alignment was adapted from previous work^11^. (**b**) Serial dilutions of cells expressing CapRel^SJ46^ or a FLAG-tagged version from its native promoter and Gp54^Bas11^ or a HA-tagged version from an arabinose-inducible promoter on media containing glucose or arabinose. (**c**) Summary of 3 independent replicates of phage spotting assay in Fig. 2d. Asterisks indicate p < 0.05 (unpaired two-tailed t-test). (**d**) Serial dilutions of phage T7 spotted on lawns of cells harboring an empty vector (EV) or a plasmid expressing CapRel^SJ46^, CapRel^Ebc^ or the chimera. Data also shown in previous work^11^.

**Figure S3.**
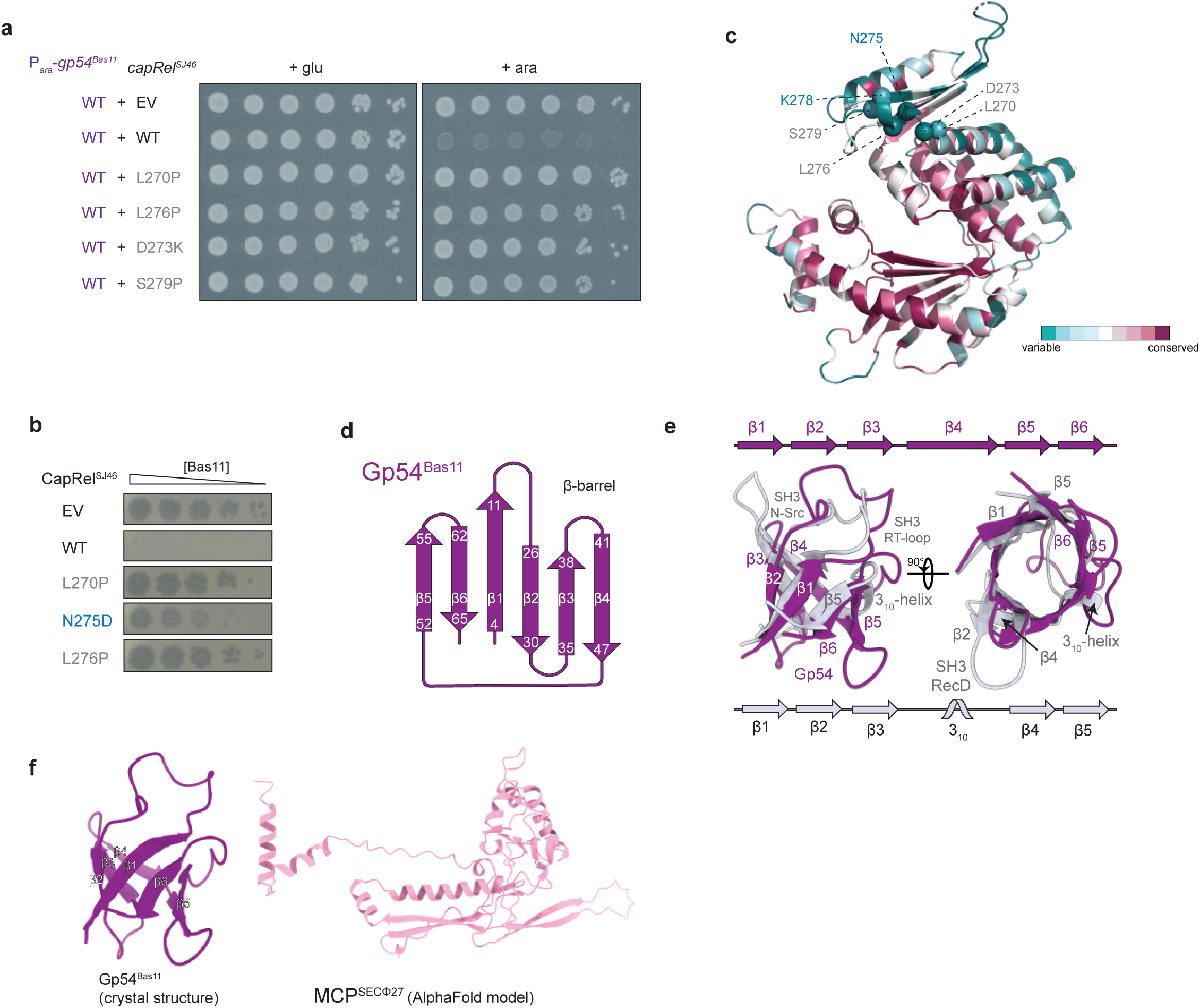
Structural analysis of Gp54^Bas11^ and its interaction with CapRel^SJ46^. (**a**) Serial dilutions of cells expressing the indicated variant of CapRel^SJ46^ from its native promoter and the wild-type Gp54^Bas11^ from an arabinose-inducible promoter on media containing glucose or arabinose. (**b**) Serial dilutions of phage Bas11 spotted on lawns of cells harboring an empty vector (EV) or a plasmid expressing CapRel^SJ46^ or the indicated variant. (**c**) Cartoon representation of the crystal structure of CapRel^SJ46^ colored by the conservation score of each amino acid calculated by ConSurf^26^. Substitutions in the α-helix (α9) formed by residues 270-279, are labeled as spheres. (**d**) Topological representation of Gp54^Bas11^. (**e**) Comparison of Gp54^Bas11^ with the 5 β-stranded β-barrel SH3 domain (residues 470 to 533) from the RecD subunit of the RecBCD repair complex (PDB 5LD2^27^). The two domains superimposed with a Z-score of 3.8. The secondary structural elements of both proteins are labeled. (**f**) Cartoon representation of the crystal structure of Gp54^Bas11^ (*left*) and the AlphaFold predicted structure of MCP^SECΦ27^ (*right*).

**Figure S4.**
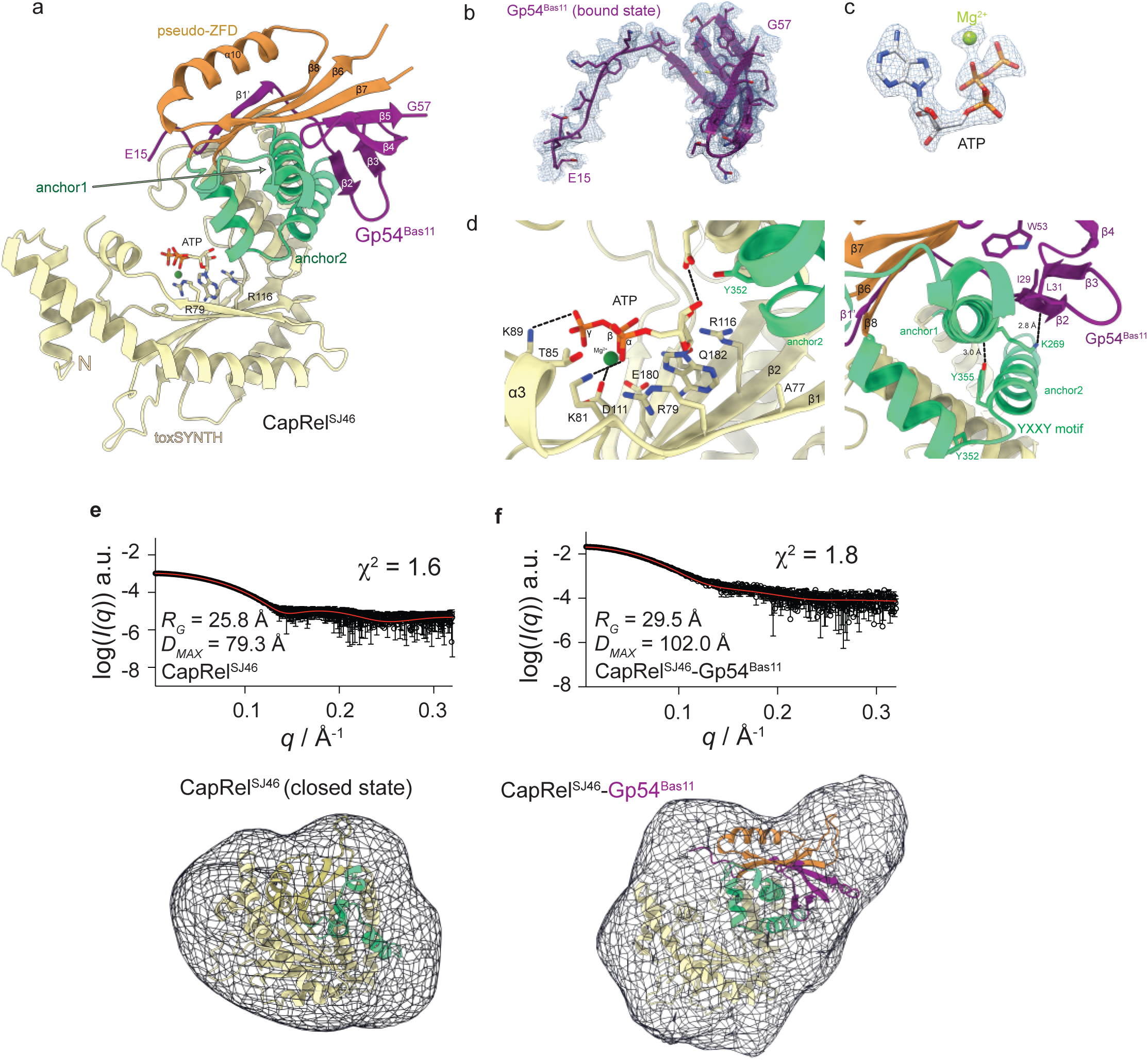
Structural analysis of the CapRel^SJ46^-Gp54^Bas11^ complex. (a) Cartoon representation of the crystal structure of CapRel^SJ46^-Gp54^Bas11^ complex bound to ATP. ATP coordination residues of CapRel^SJ46^ toxin domain (R79 and R116) are labeled. (b) Cartoon representation and the corresponding m2Fo-DFc electron density map of Gp54^Bas11^ in the CapRel^SJ46^-bound state. (c) The unbiased mFo-DFc electron density map of ATP and Mg^2+^ observed in the crystal structure of CapRel^SJ46^-Gp54^Bas11^ complex. (d) Details interface of CapRel^SJ46^-Gp54^Bas11^ complex structure indicating interaction with Gp54^Bas11^ via K269 (*right*) tethers Y355 of the YXXY neutralization motif away from the toxin active site of CapRel^SJ46^ (*left)*. (e) *Top*, experimental SAXS analysis of CapRel^SJ46^ (open black circles). The theoretical scattering of CapRel^SJ46^ in the closed state predicted by AlphaFold is shown in red. *Bottom,* comparison of the model of CapRel^SJ46^ in the unbound closed state with an ab-initio envelope calculated from the experimental SAXS data using DAMMIF^28^. (f) *Top,* experimental SAXS analysis of CapRel^SJ46^-Gp54^Bas11^ complex (open black circles). The theoretical scattering of the CapRel^SJ46^-Gp54^Bas11^ crystal structure is shown in red. *Bottom*, comparison of the crystal structure of CapRel^SJ46^-Gp54^Bas11^ with an ab-initio envelope calculated from the experimental SAXS data using DAMMIF^28^.

**Figure S5.**
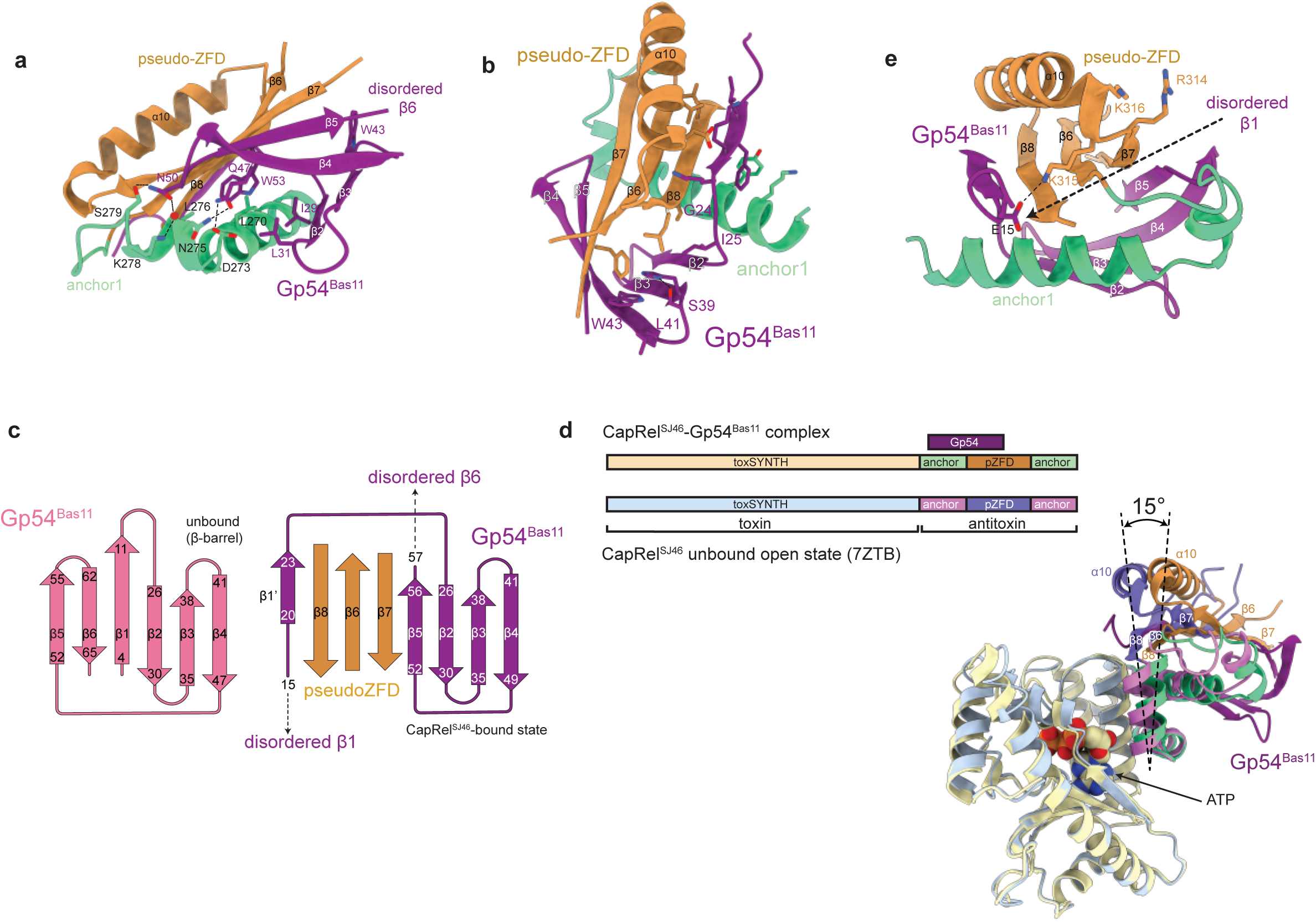
Gp54^Bas11^ interacts with the antitoxin region of CapRel^SJ46^. (**a**) Details of the interface formed by the antitoxin domain of CapRel^SJ46^ and Gp54^Bas11^ observed in the crystal structure. Residues in anchor-1 of CapRel^SJ46^ and residues in Gp54^Bas11^ involved in the interface are labeled. (**b**) Detailed interface of Gp54^Bas11^ bound to CapRel^SJ46^. Residues in Gp54^Bas11^ that are involved in hydrophobic interaction with CapRel^SJ46^ are labeled. (**c**) Topological representation of Gp54^Bas11^ in an unbound, β-barrel state (*left*) or a CapRel^SJ46^-bound state involving interaction with the pseudo-ZFD of CapRel^SJ46^ (*right*). (**d**) Superposition and comparison of CapRel^SJ46^ in complex with Gp54^Bas11^ and unbound open state (PDB 7ZTB). (**e**) Details of the interface of CapRel^SJ46^ bound to Gp54^Bas11^, with residues 314-316 of CapRel^SJ46^ labeled. The most N-terminal residue observed in crystal structure (E15) of Gp54^Bas11^ and the disordered β1 are indicated.

**Figure S6.**
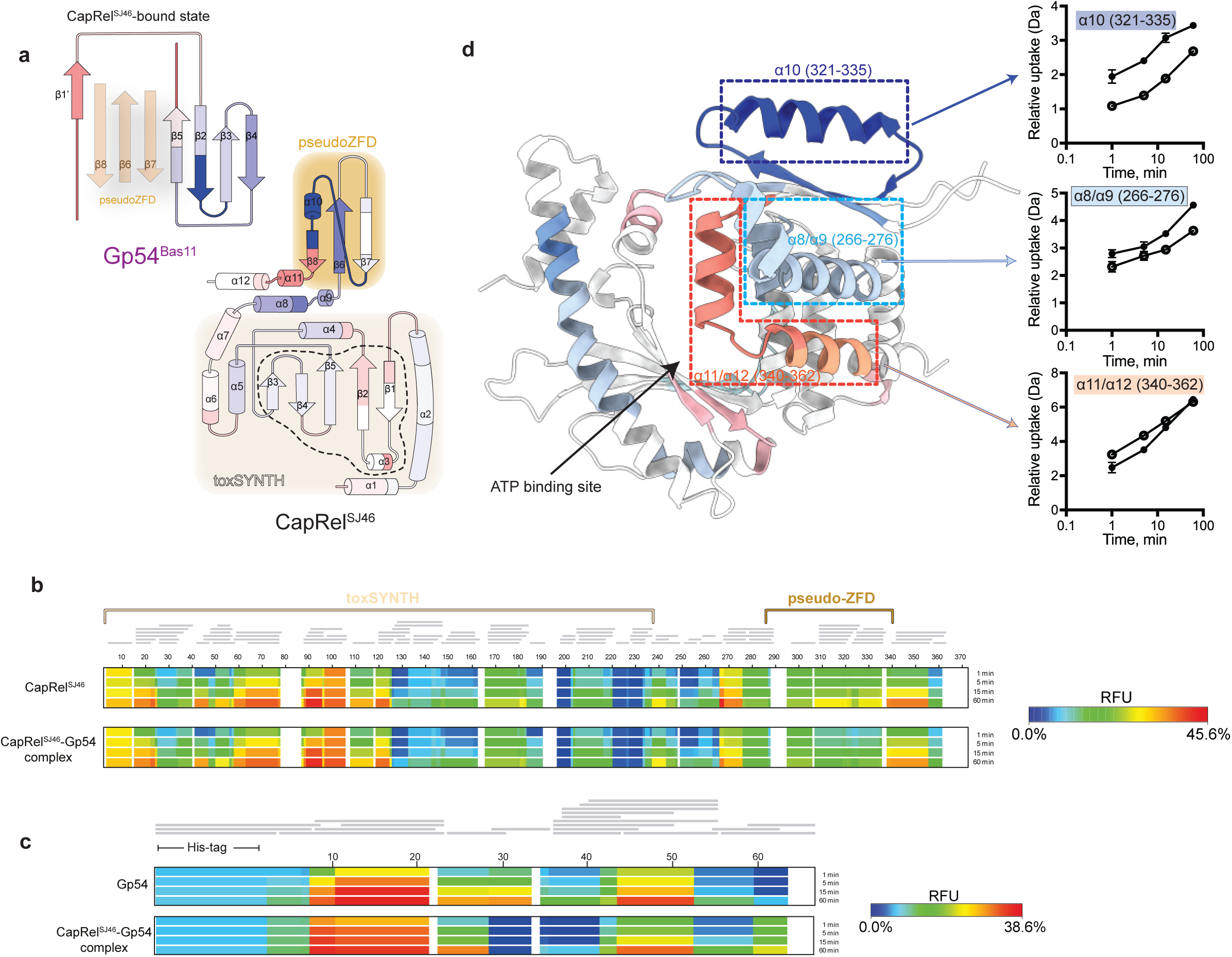
Characterization of the CapRel^SJ46^-Gp54^Bas11^ and CapRel^SJ46^-MCP^SECΦ27^ interactions. (**a**) Topological representation of CapRel^SJ46^ and Gp54^Bas11^ colored according to the ΔHDX as in Fig. 3e-f. (**b**) Heat maps representing the HDX of CapRel^SJ46^ (*top*) and CapRel^SJ46^-Gp54^Bas11^ complex (*bottom*). The relative fractional uptake (RFU) is indicated based on the color scale shown. (**c**) Heat maps representing the HDX of Gp54^Bas11^ (*top*) and CapRel^SJ46^-Gp54^Bas11^ complex (*bottom*). The relative fractional uptake (RFU) is indicated based on the color scale shown. (**d**) *Left*, ΔHDX after 5 min, between CapRel^SJ46^ and the CapRel^SJ46^-Gp54^Bas11^ complex plotted as a heat map on the structure of CapRel^SJ46^ in the open, active state. *Right*, evolution of the deuterium uptake kinetics of peptides from α8/α9 (266-276), α10 (321-335), and α11/α12 (340-362) of CapRel^SJ46^ in the unbound (solid black circles) and Gp54^Bas11^-bound (open black circles) states.

**Figure S7.**
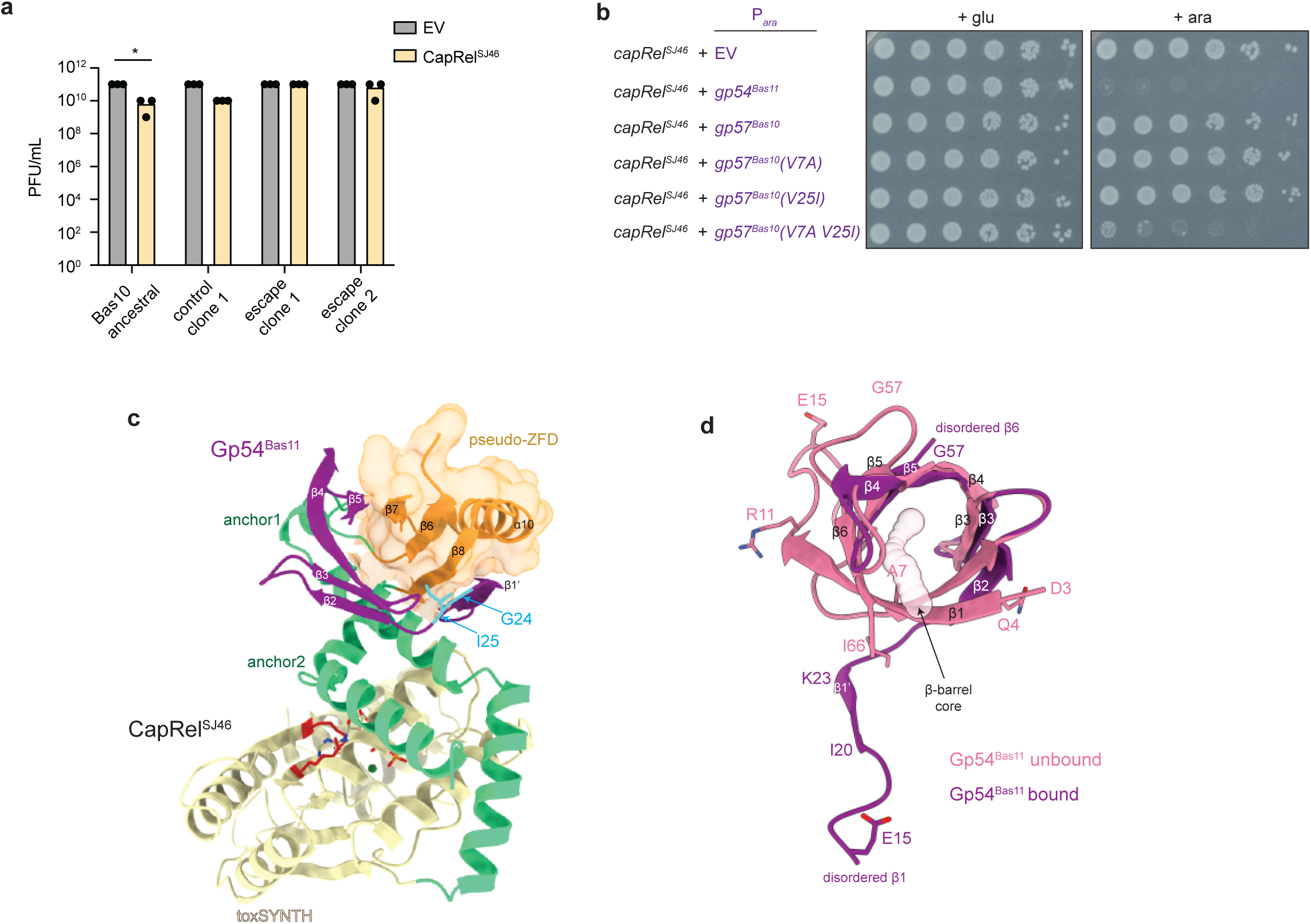
Analysis of Gp54 homologs in related phages. (**a**) Summary of 3 independent replicates of phage spotting assay in Fig. 4b. Asterisks indicate p < 0.05 (unpaired two-tailed t-test). (**b**) Serial dilutions of cells expressing CapRel^SJ46^ from its native promoter and Gp54^Bas11^ or Gp57^Bas10^ (the Gp54 homolog in Bas10) or the indicated variant from an arabinose-inducible promoter on media containing glucose or arabinose. (**c**) The crystal structure of Gp54^Bas11^ (purple) bound to CapRel^SJ46^ (colored by domains). Substitutions in Gp54^Bas11^ observed in Bas11 escape mutant (G24D) and residue I25 that is different between Gp54^Bas11^ and Gp57^Bas10^ are colored in cyan. (**d**) Comparison of the unbound (pink) and the CapRel^SJ46^-bound state (purple) of Gp54^Bas11^, indicating the structural changes. Residue A7V might stabilize the β-barrel core in the unbound state.

**Figure S8.**
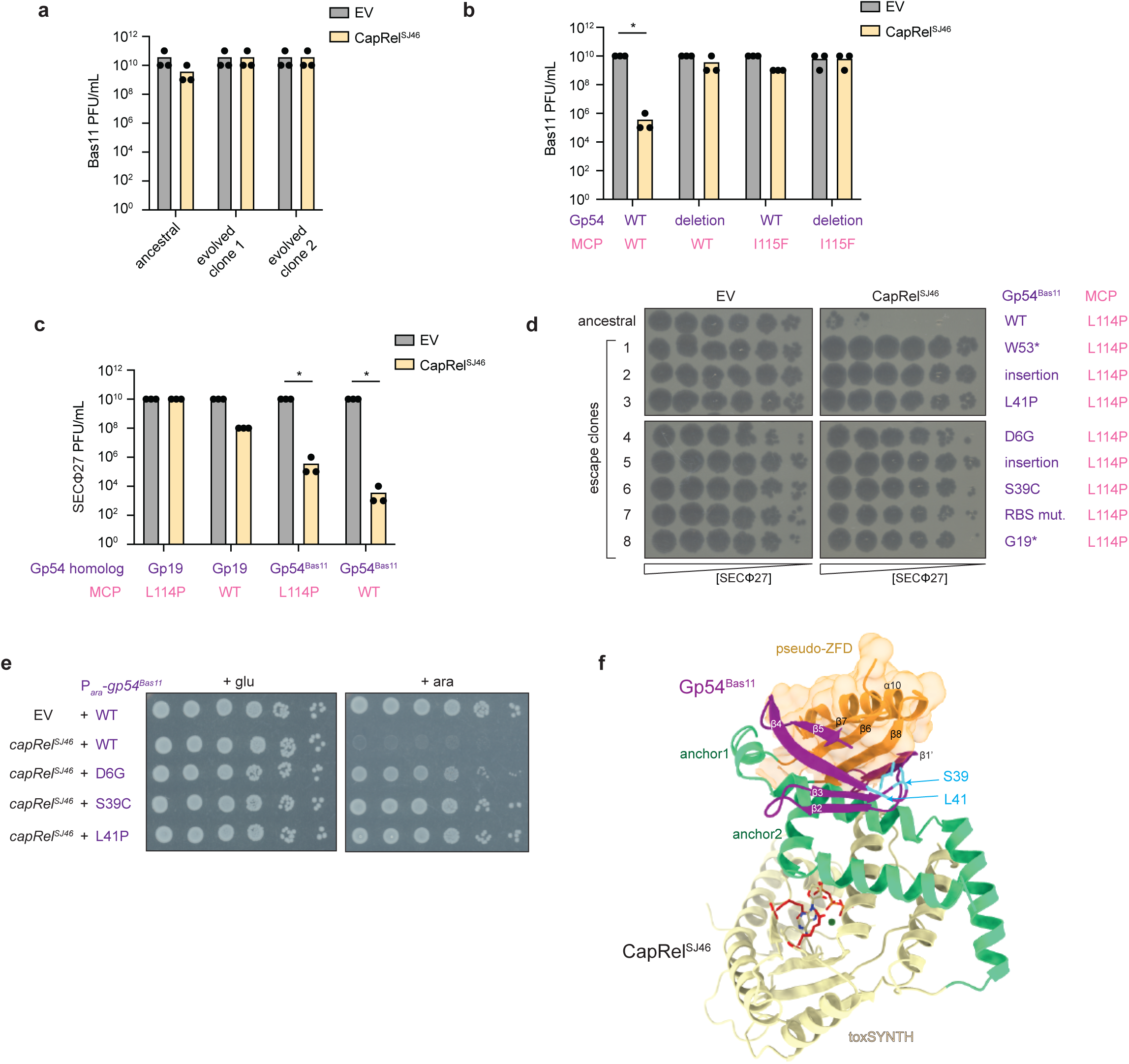
Both Gp54^Bas11^ and MCP activate CapRel^SJ46^ in Bas11 phage. (**a**) Summary of 3 independent replicates of phage spotting assay in Fig. 5a. (**b**) Summary of 3 independent replicates of phage spotting assay in Fig. 5b. Asterisks indicate p < 0.05 (unpaired two-tailed t-test). (**c**) Summary of 3 independent replicates of phage spotting assay in Fig. 5c. Asterisks indicate p < 0.05 (unpaired two-tailed t-test). (**d**) Serial dilutions of the ancestral SECΦ27 MCP(L114P) phage harboring Gp54^Bas11^ and eight escape clones spotted on lawns of cells harboring an empty vector (EV) or a plasmid expressing CapRel^SJ46^. The corresponding genotypes of its MCP or Gp54^Bas11^ were indicated on the right. Three independent replicates are shown in Fig. 5d. (**e**) Serial dilutions of cells expressing CapRel^SJ46^ from its native promoter and the indicated variant of Gp54^Bas11^ from an arabinose-inducible promoter on media containing glucose or arabinose. (**f**) The crystal structure of Gp54^Bas11^ (purple) bound to CapRel^SJ46^ (colored by domains). Residues S39 and L41 in Gp54^Bas11^ substituted in the SECΦ27 MCP(L114P) escape mutants are colored in cyan.

## Methods

### Strains and growth conditions

All bacterial and phage strains used in this study are listed in Table S3. *Escherichia coli* strains were routinely grown at 37 °C in Luria broth (LB) medium for cloning and maintenance. Phages were propagated by infecting a culture of *E. coli* MG1655 at an OD_600_ ∼0.1-0.2 with a MOI of 0.1. Cleared cultures were pelleted by centrifugation to remove residual bacteria and filtered through a 0.2 μm filter. Chloroform was then added to phage lysates to prevent bacterial growth. All phage infection experiments were performed in LB medium at 25 °C. Antibiotics were used at the following concentrations (liquid; plates): carbenicillin (50 μg/mL; 100 μg/mL), chloramphenicol (20 μg/mL; 30 μg/mL).

### Plasmid construction

All plasmids are listed in Table S4. All primers are listed in Table S5.

pBAD33-*gp54^Bas11^* constructs: wild-type or mutant variants of *gp54^Bas11^* were PCR-amplified from the corresponding wild-type Bas11 or escaping phage clones using primers TZ-3 and TZ-4, and inserted into pBAD33 linearized with TZ-1 and TZ-2 using Gibson assembly. To add a C-terminal HA-tag, primers TZ-9 and TZ-10 were used to PCR-amplify pBAD33-*gp54^Bas11^* followed by Gibson assembly. Mutations that produce the single amino-acid substitutions D6E, A7V, and I25V in Gp54^Bas11^ were generated by site-directed mutagenesis using primers TZ-38 to TZ-43.

pBR322-*capRel^SJ46^* constructs: mutations that produce the single amino-acid substitutions N275D, D273K, K278E, S279P were generated by site-directed mutagenesis using primers TZ-11 to TZ-18. Mutations that produce the triple substitution K278E R314E K316E were introduced by two-step site-directed mutagenesis using primers TZ-23 and TZ-24, then TZ-15 and TZ-16. To add a C-terminal FLAG-tag to CapRel^SJ46^, primers TZ-25 and TZ-26 were used to PCR-amplify pBR322-*capRel^SJ46^*followed by Gibson assembly.

pBAD33*-gp54^Bas11^ homolog* constructs: the genes encoding Gp54^Bas11^ homologs (Gp57^Bas10^, Gp60^Bas8^, Gp57^Bas5^ and Gp19^SECΦ27^) were PCR-amplified from the corresponding phage using primers TZ-30 to TZ-37 and inserted into linearized pBAD33 by Gibson assembly. Mutations that produce the single amino-acid substitutions V7A and V25I in Gp57^Bas10^ were generated by site-directed mutagenesis using primers TZ-44 to TZ-47.

pET-*His_6_-gp54^Bas11^*constructs: wild-type or the G24D variant of *gp54^Bas11^* was PCR-amplified from the corresponding phage using primers TZ-7 and TZ-8, and inserted into pET-His_6_ vector linearized with primers TZ-5 and TZ-6 using Gibson assembly.

pET-*His_6_-MBP-capRel^SJ46^* construct: *capRel^SJ46^*was first PCR-amplified from pBR322-*capRel^SJ46^* using primers TZ-48 and TZ-49, and inserted into pET-His_6_ vector linearized with primers TZ-5 and TZ-6 using Gibson assembly. The gene encoding MBP was PCR-amplified with TZ-52 and TZ-53, and inserted into pET-*His_6_-capRel^SJ46^* linearized with primer TZ-50 and TZ-51 using Gibson assembly.

pBAD33-*mcp^Bas10^*construct: the gene encoding the major capsid protein of Bas10 (Gp9^Bas10^) was PCR-amplified from phage Bas10 using primers TZ-27 and TZ-28, and inserted into pBAD33 linearized with primers TZ-29 and TZ-1 using Gibson assembly.

### Strain construction

Plasmids described above were introduced into *E. coli* MG1655 by TSS transformation or electroporation.

Bas11 mutant phage producing MCP(I115F) were generated using a CRISPR-Cas system for targeted mutagenesis as described previously^29^. Briefly, sequences for RNA guides to target Cas9-mediated cleavage were designed using the toolbox in Geneious Prime 2022.0.2 and selected for targeting of *mcp^Bas11^* (Gp8 in Bas11) but nowhere else in the Bas11 genome. The guides were inserted into the pCas9 plasmid and tested for their ability to restrict Bas11. An efficient guide was selected and the pCas9-*guide* plasmid was co-transformed into *E. coli* MG1655 with a high copy-number repair plasmid containing *mcp^Bas11^(I115F)* with the guide mutated synonymously to prevent self-cutting. The wild-type Bas11 phage was plated onto a strain containing both the pCas9-*guide* and the repair plasmid, and single plaques were screened by Sanger Sequencing. Two clones that produce the I115F substituted MCP^Bas11^ were propagated on strains containing only pCas9-*guide* for further selection.

SECΦ27 mutant phages producing Gp54^Bas11^ instead of its homolog in SECΦ27 (Gp19) were similarly generated as described above. The guide was selected such that it only targeted gene *19* in SECΦ27, but not gene *54* from Bas11. The selected pCas9-*guide* plasmid was co-transformed into *E. coli* MG1655 with a high copy-number repair plasmid containing the coding sequence of gene *54* from Bas11 flanked by the region that flanks gene *19* in SECΦ27 for homologous recombination. The wild-type SECΦ27 phage or the mutant producing MCP(L114P) was plated onto the strain containing pCas9 plasmid and the repair plasmid for selection. Two clones each were propagated and selected twice on strains containing only pCas9-*guide*.

### Phage spotting assays and efficiency of plaquing (EOP) measurements

Phage spotting assays were conducted similarly to that described previously^11^. Phage stocks isolated from single plaques were propagated in *E. coli* MG1655 at 37 °C in LB. To titer phage, dilutions of stocks were mixed with *E. coli* MG1655 and melted LB + 0.5% agar and spread on LB + 1.2% agar plates and incubated at 37 °C overnight. For phage spotting assays, 80 μL of a bacterial strain of interest was mixed with 4 mL LB + 0.5% agar and spread on an LB + 1.2% agar + antibiotic plate. Phage stocks were then serially diluted in 1x FM buffer (20 mM Tris-HCl pH 7.4, 100 mM NaCl, 10 mM MgSO_4_), and 2 μL of each dilution was spotted on the bacterial lawn. Plates were then incubated at 25 °C overnight before imaging. Efficiency of plaquing (EOP) was calculated by comparing the ability of the phage to form plaques on an experimental strain relative to the control strain. Experiments were replicated 3 times independently and representative images are shown.

### Toxicity assays on solid media

Bacterial toxicity assays were conducted similarly to that described previously^11^. For co-producing CapRel^SJ46^ with Gp54 homologs, or with major capsid proteins, single colonies of *E. coli* MG1655 harboring pBR322-*capRel^SJ46^* and pBAD33-*Gp54 homolog* or pBAD33-*capsid protein* (wild-type or the corresponding variants) were grown for 6 hours at 37 °C in LB-glucose to saturation. 200 μL of each saturated culture was then pelleted by centrifugation at 4000 *g* for 10 min, washed once in 1x phosphate-buffered saline (PBS), and resuspended in 400 μL 1x PBS. Cultures were then serially-diluted 10-fold in 1x PBS and spotted on M9L plates supplemented with 0.4% glucose or 0.2% arabinose. M9L plates contain M9 medium (6.4 g/L Na_2_HPO_4_-7H_2_O, 1.5 g/L KH_2_PO_4_, 0.25 g/L NaCl, 0.5 g/L NH_4_Cl medium supplemented with 0.1% casamino acids, 0.4% glycerol, 2 mM MgSO_4_, and 0.1 mM CaCl_2_) supplemented with 5% LB (v/v). Plates were then incubated at 37 °C overnight before imaging.

### Isolation of phage escape mutants to infect CapRel^SJ46^

Bas11 or SECΦ27 MCP(L114P) escape mutants were isolated by plating a population of phage onto CapRel^SJ46^-containing cells. 20 µL of 10^10^ pfu/mL Bas11 or SECΦ27 MCP(L114P) phage mixed with 40 µL overnight culture of *E. coli* MG1655 pBR322-*capRel^SJ46^* were added to 4 mL of LB + 0.5% agar and spread onto LB + 1.2% agar. Plates were incubated at 25 °C overnight. Single plaques were isolated and propagated using the same strain in LB at 25 °C. Amplified phage lysates were pelleted to remove bacteria, and sequenced by Illumina sequencing as described below to identify mutations.

Bas10 or Bas11 phage with gene *54* deleted were evolved to completely overcome CapRel^SJ46^ defense using an experimental evolution protocol as described previously^30^. Briefly, five independent populations were evolved in a 96-well plate containing a sensitive host *E. coli* MG1655 pBR322-EV and a resistant host *E. coli* MG1655 pBR322-*capRel^SJ46^*. One control population was evolved with only the sensitive host. Overnight bacterial cultures were back-diluted to OD_600_ = 0.01 in LB and 100 μL was seeded into each well. Cells were infected with 10-fold serial dilutions of Bas10 or Bas11 phage with gene *54* deleted with MOI from 100 to 10^-4^, with one well uninfected to monitor for contamination. Plates were sealed with breathable plate seals and incubated at 25 °C for 14 hours (for Bas10) or 17 hours (for Bas11) in a plate shaker at 1000 rpm. Cleared wells from each population were pooled, pelleted at 4000 *g* for 20 min to remove bacteria, and the supernatant lysates were transferred to a 96 deep-well block with 40 µL chloroform added to prevent bacterial growth. Lysates were spotted onto both sensitive and resistant hosts to check the defense phenotype. Three rounds of evolution were performed for Bas10 and four populations were able to overcome CapRel^SJ46^ defense. Two rounds of evolution were performed for Bas11 phage with gene *54* deleted. Evolved clones from each evolved population were isolated by plating to single plaques on lawns of resistant host, and control clones from the control population were isolated on a lawn of the sensitive host. Two clones from each population were propagated using the corresponding host and sequenced as described below.

### Phage DNA extraction and Illumina sequencing

Phage DNA extraction and sequencing were conducted as described previously^11^. To extract phage DNA, high titer phage lysates (> 10^6^ pfu/µL) were treated with DNase I (0.001 U/µL) and RNase A (0.05 mg/mL) at 37 °C for 30 min. 10 mM EDTA was used to inactivate the nucleases. Lysates were then incubated with Proteinase K at 50 °C for 30 min to disrupt capsids and release phage DNA. Phage DNA was isolated by ethanol precipitation. Briefly, NaOAc pH 5.2 was added to 300 mM followed by 100% ethanol to a final volume fraction of 70%. Samples were incubated at -80 °C overnight, pelleted at 21,000 *g* for 20 min and supernatant removed. Pellets were washed with 100 µL isopropanol and 200 µL 70% (v/v) ethanol, and then aired dried at room temperature and resuspended in 25 µL 1x TE buffer (10 mM Tris-HCl, 0.1 mM EDTA, pH = 8). Concentrations of extracted DNA were measured by NanoDrop (Thermo Fisher Scientific).

To prepare Illumina sequencing libraries, 100-200 ng of genomic DNA was sheared in a Diagenode Bioruptor 300 sonicator water bath for 20x 30 s cycles at maximum intensity. Sheared genomic DNA was purified using AmpureXP beads, followed by end repair, 3’ adenylation, and adaptor ligation. Barcodes were added to both 5’ and 3’ ends by PCR with primers that anneal to the Illumina adaptors. The libraries were cleaned by Ampure XP beads using a double cut to elute fragment sizes matching the read-lengths of the sequencing run. Libraries were sequenced on an Illumina MiSeq at the MIT BioMicro Center. Illumina reads were assembled to the reference genomes using Geneious Prime 2022.0.2.

### Co-immunoprecipitation (co-IP) analysis

Co-immunoprecipitation experiments were conducted similar to those described previously^11^. For co-producing CapRel^SJ46^ and Gp54^Bas11^ or with MCP ^SECΦ27^, *E. coli* MG1655 containing pBR322-*capRel^SJ46^*or pBR322-*capRel^SJ46^-FLAG* (wild type or mutant variants) and pBAD33-*gp54^Bas11^-HA* (wild type or mutant variants) or pBAD33-*mcp^SECΦ27^*-HA were grown overnight in M9-glucose. Overnight cultures were back-diluted to OD_600_ = 0.05 in 50 mL of M9 (no glucose) and grown to OD_600_ ∼ 0.3 at 37 °C. Cells were induced with 0.2% arabinose for 30 min at 37 °C, then OD_600_ was measured and cells were pelleted at 4000 *g* for 10 min at 4 °C. Supernatant was removed and cells were resuspended in 800 μL lysis buffer (25 mM Tris-HCl, 150 mM NaCl, 1 mM EDTA, 1% Triton X-100 and 5% glycerol) supplemented with protease inhibitor (Roche), 1 μL/mL Ready-Lyse^TM^ Lysozyme Solution (Lucigen) and 1 μL/mL benzonase nuclease (Sigma). Samples were lysed by two freeze-thaw cycles, and lysates were normalized by OD_600_. Lysates were pelleted at 21,000 *g* for 10 min at 4 °C, and 750 μL of supernatant were incubated with pre-washed anti-FLAG magnetic agarose beads (Pierce) for 1 hour at 4 °C with end-over-end rotation. Beads were then washed 3 times with 500 μL lysis buffer. 1x Laemmli sample buffer (Bio-Rad) supplemented with 2-mercaptoethanol was added to beads directly to elute proteins.

Samples were boiled at 95 °C and analyzed by 4-20% SDS-PAGE and transferred to a 0.2 μm PVDF membrane. Anti-FLAG and anti-HA antibodies (Cell Signaling Technology) were used at a final concentration of 1:1000, and SuperSignal West Femto Maximum Sensitivity Substrate (ThermoFisher) was used to develop the blots. Blots were imaged by a ChemiDoc Imaging system (Bio-Rad). Images shown are representatives of two independent biological replicates.

### Error-prone PCR mutagenesis of CapRel^SJ46^ and selection with Gp54^Bas11^

The C-terminus of CapRel^SJ46^ was mutagenized using error-prone PCR based mutagenesis as described previously^11^. Briefly, primers TZ-54 and TZ-55 were used to amplify the C-terminus of CapRel^SJ46^ using Taq polymerase (NEB) and 0.5 mM MnCl_2_ was added to the reaction as the mutagenic agent. PCR products were treated with Dpn I, column purified, and inserted into pBR322-*capRel^SJ46^* backbone amplified with primer TZ-56 and TZ-57 using Gibson assembly. Gibson products were transformed into DH5α and grown overnight in LB at 37 °C. Overnight cultures were miniprepped to obtain the mutagenized library. Individual colonies were Sanger sequenced to assess the number of mutations. To perform the selection, mutagenized library was electroporated into *E. coli* MG1655 pBAD33-*gp54^Bas11^*, and plated onto M9L plates containing 0.2% arabinose to select for survivors. Colonies were picked and sequenced to identify mutations in CapRel^SJ46^.

### Protein expression and purifications

To produce His_6_-MBP-tagged CapRel^SJ46^, *E. coli* BL21(DE3) cells were transformed with pET-*His_6_-MBP-capRel^SJ46^* and grown in LB medium to OD_600_ of 0.5. Protein expression was induced by addition of 0.3 mM IPTG, and cells were grown for 3 hours at 30 °C. The culture was centrifuged at 4000 *g* for 10 min at 4 °C, and cell pellet was resuspended in lysis buffer (50 mM Tris-HCl pH 8.0, 500 mM NaCl, 500 mM KCl, 2 mM MgCl_2_, 1 mM DTT) supplemented with 0.4 mM PMSF, 10 μg/mL lysozyme and 7.5 U/mL benzonase nuclease (Milipore). Cells were disrupted using sonication (Qsonica) and glycerol was added to the lysate at final 10% concentration after sonication. The supernatant was separated from the pellet by centrifugation (15,000 rpm for 30 min, JA-25.50 rotor (Beckman Coulter)). The clarified supernatant was loaded onto a gravity-flow chromatography column (BioRad) packed with 2 mL Ni-NTA agarose resin (Qiagen) pre-equilibrated with 15 mL lysis buffer. The resin was washed with 10 column volumes of wash buffer 1 (50 mM Tris-HCl pH 8.0, 500 mM NaCl, 500 mM KCl, 2 mM MgCl_2_, 10 mM imidazole, 10% glycerol, 1 mM DTT), and then with 10 column volumes of wash buffer 2 (50 mM Tris-HCl pH 8.0, 500 mM NaCl, 500 mM KCl, 2 mM MgCl_2_, 50 mM imidazole, 10% glycerol, 1 mM DTT). The proteins were eluted in 4 mL elution buffer (50 mM Tris-HCl pH 8.0, 500 mM NaCl, 500 mM KCl, 2 mM MgCl_2_, 300 mM imidazole, 10% glycerol, 1 mM DTT). To remove remaining contaminants, the eluted protein sample was loaded onto a size exclusion chromatography (SEC) Superdex 200 Increase 10/300 GL column (Cytiva) pre-equilibrated in the SEC buffer (50 mM Tris-HCl pH 8.0, 250 mM NaCl, 250 mM KCl, 2 mM MgCl_2_, 1 mM DTT). The fractions containing the protein of interest were pooled and concentrated to around 1 mg/mL. Purity of the protein samples were assessed spectrophotometrically and by SDS-PAGE.

To produce His_6_-tagged Gp54^Bas11^ or the G24D variant, *E. coli* BL21(DE3) cells were transformed with pET-*His_6_-gp54^Bas11^* (wild-type or G24D) and grown in LB medium to OD_600_ of 0.5. Protein expression was induced by addition of 0.3 mM IPTG, and cells were grown for 3 hours at 30 °C. Purification steps were performed similarly as described above with the following buffers. Lysis buffer contains 50 mM Tris-HCl pH 8.0, 150 mM NaCl, 2 mM MgCl_2_, 1 mM DTT supplemented with 0.4 mM PMSF, 10 μg/mL lysozyme and 7.5 U/mL benzonase nuclease (Milipore). Wash buffer 1 contains 50 mM Tris-HCl pH 8.0, 500 mM NaCl, 2 mM MgCl_2_, 10 mM imidazole, 10% glycerol, 1 mM DTT. Wash buffer 2 contains 50 mM Tris-HCl pH 8.0, 150 mM NaCl, 2 mM MgCl_2_, 50 mM imidazole, 10% glycerol, 1 mM DTT. Elution buffer contains 50 mM Tris-HCl pH 8.0, 150 mM NaCl, 2 mM MgCl_2_, 300 mM imidazole, 10% glycerol, 1 mM DTT. To remove remaining contaminants, the eluted protein sample was loaded onto a size exclusion chromatography (SEC) Superose 6 Increase 10/300 GL column (Cytiva) pre-equilibrated in the SEC buffer (50 mM Tris-HCl pH 8.0, 150 mM NaCl, 2 mM MgCl_2_, 1 mM DTT). The fractions containing the protein of interest were pooled and concentrated to around 5 mg/mL. Purity of the protein samples were assessed spectrophotometrically and by SDS-PAGE.

### Cell-free translation

Experiments with PURExpress *in vitro* protein synthesis kit (NEB, E6800) were performed as per the manufacturer’s instructions. All reactions were supplemented with 0.8 U/µL RNase Inhibitor Murine (NEB, M0314S). Purified His_6_-MBP-tagged CapRel^SJ46^ protein was added to the reaction at a final concentration of 500 nM, and purified His_6_-tagged Gp54^Bas11^ or the G24D variant was used at a final concentration of 4 µM. A template plasmid encoding the control protein DHFR (provided by the kit) was used at 6 ng/µL. The reactions were incubated at 37 °C for 2 hours, and 2 µL of each reaction was mixed with 10 µL of 1x Laemmli sample buffer (Bio-Rad) supplemented with 2-mercaptoethanol. The mixtures were boiled for 5 min at 95 °C and analyzed by 12% SDS-PAGE. The gels were stained with Coomassie stain and imaged by a ChemiDoc Imaging system (Bio-Rad). Images shown are representatives of three independent biological replicates.

### Crystallization and structure determination of Gp54^Bas11^ and the CapRel^SJ46^-Gp54^Bas11^ complex

His_6_-tagged Gp54^Bas11^ was purified as described above and concentrated to 5 mg/mL for crystallization. Crystallization conditions for His_6_-Gp54^Bas11^ were screened by sitting drop vapor diffusion using a Formulatrix NT8 drop setter and commercial screening kits. Each drop, consisting of 100 nL of protein solution plus 100 nL of reservoir solution, was equilibrated against 70 µL of reservoir solution. Crystals appeared in Index HT (Hampton Research) condition B12 (2.8 M sodium acetate trihydrate pH 7). These conditions were optimized, and the final crystals were grown by hanging drop vapor diffusion, with drops consisting of 2 µL protein plus 2 µL of well solution (3.2 M potassium acetate pH 7.0) at room temperature. After 8 days, a crystal was harvested and directly vitrified in a nitrogen gas stream at 100 K (Oxford Cryostream). X-ray diffraction data was collected on a Rigaku Micromax-007 rotating anode with Osmic VariMax-HF mirrors and a Rigaku Saturn 944 detector. Diffraction data were processed with the XDS suite^31^. Phaser^32^ was used to solve the structure by molecular replacement using an AlphaFold^33^ model. The molecular replacement solution was refined in PHENIX^34^ with manual model building done with Coot^35^. The model was refined to final R_work_/R_free_ of 0.211/0.252. The X-ray data collection and refinement statistics are summarized in Table S1.

For the CapRel^SJ46^-Gp54^Bas11^ complex, CapRel^SJ46^ and Gp54^Bas11^ were purified as described above, and mixed in a 1:1 ratio at a concentration of 2 mg/ml. The complex was then further purified by SEC (in (50 mM Tris-HCl pH 8.0, 150 mM NaCl, 2 mM MgCl_2_, 1 mM DTT)) and the resulting sample concentrated to 10 mg/mL for crystallization. Crystallization conditions for the CapRel^SJ46^-Gp54^Bas11^ complex were screened as such or supplemented with 5 mM of ATP. Crystals grew within a week in 25% PEG 1500 in a malic acid, MES, Tris buffer system (pH 8.0). Prior to data collection the crystals were cryo-protected by soaking in the mother liquor solution supplemented with 25% glycerol, and flash frozen in liquid nitrogen for storage. X-ray diffraction data was collected at the I24 beamline of the Diamond Light Source synchrotron (UK) on a CdTe Eiger2 9M detector. The data were processed using XDS suite^31^ and scaled with Aimless. The structure was solved by molecular replacement performed with Phaser^32^ using the coordinates of the toxSYNTH domain of CapRel^SJ46^ (PDB ID: 7ZTB). Initial automated model building was performed with Buccaneer^36^ which partially completed Gp54^Bas11^ and further improved with the MR-Rosetta suite from the Phenix package^37^. After several iterations of manual building with Coot^35^ and maximum likelihood refinement as implemented in Buster/TNT^38^, the model was refined to R/Rfree of 0.193/0.236. The X-ray data collection and refinement statistics are summarized in Table S1.

### Homology search, alignment, and conservation analysis

CapRel^SJ46^ homologs were identified, aligned, and used as input for ConSurf analysis as described previously^11^. Homologs of the major capsid proteins or Gp54^Bas11^ in BASEL phages were identified by BLASTp^39^ searches against each phage genome, and aligned by MUSCLE^40^.

### Hydrogen deuterium exchange mass spectrometry (HDX-MS)

HDX-MS experiments were performed on an HDX platform composed of a Synapt G2 mass spectrometer (Waters Corporation) connected to a nanoAcquity UPLC system following the protocol previously described^11^. Samples of CapRel^SJ46^, Gp54^Bas11^ and the CapRel^SJ46^-Gp54^Bas11^ were prepared at a concentration of 100 µM (the integrity of the complex was confirmed by SEC prior to the HDX-MS experiment). For each experiment 8 µL of sample were incubated for 1 min, 5 min, 15 min and 60 min in 72 µL of labelling buffer L (50 mM HEPES, 500 mM KCl, 500 mM NaCl, 2 mM MgCl2, 1 mM TCEP, 0.002% mellitic acid, pH 7.5) at 20°C. The non-deuterated reference points were prepared by replacing buffer L by equilibration buffer E (50 mM HEPES, 500 mM KCl, 500 mM NaCl, 2 mM MgCl2, 1 mM TCEP, 0.002% mellitic acid, pH 7.5). After labelling, the samples are quenched by mixing with 80 µL of pre-chilled quench buffer Q (50 mM KPhos, 1 mM TCEP, 1.2 % formic acid, pH 2.4). Then samples were directly flash-frozen in liquid nitrogen and stocked at -80°C freezer until injection. For the injection, samples were thaw at room temperature and 150 µL of the quench samples are directly transferred to the Enzymate BEH Pepsin Column (Waters Corporation) at 200 µL/min and at 20 °C with a pressure 3 kPSI. Peptic peptides were trapped for 3 min on an Acquity UPLC BEH C18 VanGuard Pre-column (Waters Corporation) at a 200 µL/min flow rate in water (0.1% formic acid in HPLC water pH 2.5) before eluted to an Acquity UPLC BEH C18 Column for chromatographic separation. Separation was done with a linear gradient buffer (3–45% gradient of 0.1% formic acid in acetonitrile) at a flow rate of 40 µL/min. Peptides identification and deuteration uptake analysis was performed on the Synapt G2 in ESI ± MSE mode (Waters Corporation). Leucine enkephalin was applied for mass accuracy correction and sodium formate was used as calibration for the mass spectrometer. MSE data were collected by a 20–30 V transfer collision energy ramp. The pepsin column was washed between injections using pepsin wash buffer (1.5 M Guanidinium HCl, 4% (v/v) acetonitrile, 0.8% (v/v) formic acid). A cleaning run was performed each three sample to prevent peptide carry-over. Optimized peptide identification and peptide coverage for all samples was performed from undeuterated controls (five replicates). All deuterium time points were performed in triplicate. The mass spectrometry proteomics data have been deposited to the ProteomeXchange Consortium via the PRIDE^41^ partner repository with the dataset identifier PXD050526.

### Small Angle X-ray Scattering (SAXS)

Samples for SAXS were concentrated to 10 mg/mL, flash frozen and stored at -80 °C. SAXS data were collected at the SWING beamline (Soleil and ESRF synchrotrons, France) on a Pilatus 2M detector using the standard beamline setup in SEC (size exclusion) mode. Samples were prepared in 500 mM NaCl, 500 mM KCl, 2 mM TCEP and 30 mM HEPES pH 7.5. SEC-SAXS was done with a Shodex KW404–4F column coupled to an HPLC system, in front of the SAXS data collection capillary. The sample flowed at 0.2 ml/min and the data collected at 10 °C. Radiation damaged frames were removed before data analysis. The data were analysed with the ATSAS suite^28^. SAXS-based models were derived from the coordinates of the X-ray structure of the CapRel^SJ46^-Gp54^Bas11^ complex and an AlphaFold model of unbound CapRel^SJ46^. The calculation of *ab initio* shapes was done with the program DAMMIF from the ATSAS package.

### Isothermal Titration Calorimetry (ITC)

All titrations were performed with an Affinity ITC (TA instruments) at 30 °C. For the titration, CapRel^SJ46^ was loaded in the instrument syringe at 200 µM and Gp54^Bas11^ was used in the cell at 10 µM. Titrations were performed in 50 mM HEPES pH 7.5, 500 mM KCl, 500 mM NaCl, 2 mM MgCl2 and 1 mM TCEP. Final concentrations were verified by the absorption using a Nanodrop One (ThermoScientific). All ITC measurements were performed by titrating 2 µl of CapRel^SJ46^ into Gp54^Bas11^ (Gp54^Bas11^(G24D) was used at 260µM) using a constant stirring rate of 75 rpm. All data were processed, buffer-corrected and analysed using the NanoAnalyse and Origin software packages.

## Acknowledgements

We thank A. Harms for sharing the BASEL phage collection, the MIT BioMicro Center and its staff for their support in sequencing, the MIT Structural Biology core and P. Rosen for help with X-ray crystallography. We thank S. Srikant, C. Vassallo, C. Beck, J. Ramseyer, and D. Saxton for comments on the manuscript and all members of the Laub lab for helpful discussions. M.T.L. is an Investigator of the Howard Hughes Medical Institute. A.G-P. was supported by the Fonds National de Recherche Scientifique (CDR J.0065.23F; PDR T.0090.22); ERC (CoG DiStRes, n° 864311), the Fonds Jean Brachet, and the Fondation Van Buuren. C.M. was supported by Fonds de la Recherche Scientifique (MIS grant F.45322.22). K.C.W. is a fellow of the FRIA. The authors acknowledge the use of the SWING beamline at the Soleil synchrotron (Gif-sur-Yvette, France) and I24 (Diamond Light Source synchrotron Oxfordshire UK).

## Author Contributions

Experiments were conceived and designed by T.Z., A.G-P., and M.T.L. All experiments were done by T.Z. except for X-ray crystallography, which was done together with D.L. and A.G-P., and ITC and HDX data which were done by A.C., A.N., K.C.W., and C.M. SAXS data were collected by A.T. Figure design, manuscript writing, and editing done by T.Z., A.G-P., and M.T.L. Project supervision and funding provided by A.G-P. and M.T.L.

## Author Information

The authors declare no competing financial interests. Correspondence and requests for materials should be addressed M.T.L.

## Data Availability

Structural data for Gp54^Bas11^ and the CapRel^SJ46^-Gp54^Bas11^ complex are available in PDB (9AXB and 9ERV, respectively). Sequencing data are available in the Sequence Read Archive (SRA) under BioProject PRJNA1084025. HDX-MS data can be accessed via ProteomeXchange with identifier PXD050526. All other data are available in the manuscript or the supplementary materials. Other previously published structures are available in PDB (7ZTB, 5LD2). UniRef90 database is publicly available. Reference phage genomes are publicly available: SECΦ27 (NC_047938.1), Bas05 (MZ501101.1), Bas08 (MZ501059.1), Bas10 (MZ501077.1), Bas11 (MZ501085.1). Materials including strains and plasmids are available upon reasonable request.

## References

1. Bernheim, A. & Sorek, R. The pan-immune system of bacteria: antiviral defence as a community resource. Nat. Rev. Microbiol. 18, 113–119 (2020).

2. Millman, A. et al. An expanded arsenal of immune systems that protect bacteria from phages. Cell Host Microbe 30, 1556–1569.e5 (2022).

3. Vassallo, C. N., Doering, C. R., Littlehale, M. L., Teodoro, G. I. C. & Laub, M. T. A functional selection reveals previously undetected anti-phage defence systems in the E. coli pangenome. Nat. Microbiol. 7, 1568–1579 (2022).

4. Rostøl, J. T. & Marraffini, L. (Ph)ighting Phages: How Bacteria Resist Their Parasites. Cell Host Microbe 25, 184–194 (2019).

5. Hampton, H. G., Watson, B. N. J. & Fineran, P. C. The arms race between bacteria and their phage foes. Nature 577, 327–336 (2020).

6. Gao, L. et al. Diverse enzymatic activities mediate antiviral immunity in prokaryotes. Science 369, 1077–1084 (2020).

7. Stokar-Avihail, A. et al. Discovery of phage determinants that confer sensitivity to bacterial immune systems. Cell S0092867423001721 (2023) doi:10.1016/j.cell.2023.02.029.

8. Gao, L. A. et al. Prokaryotic innate immunity through pattern recognition of conserved viral proteins. Science 377, eabm4096 (2022).

9. Garb, J. et al. Multiple phage resistance systems inhibit infection via SIR2-dependent NAD+ depletion. Nat. Microbiol. 7, 1849–1856 (2022).

10. Banh, D. V. et al. Bacterial cGAS senses a viral RNA to initiate immunity. Nature 623, 1001–1008 (2023).

11. Zhang, T. et al. Direct activation of a bacterial innate immune system by a viral capsid protein. Nature 612, 132–140 (2022).

12. Brubaker, S. W., Bonham, K. S., Zanoni, I. & Kagan, J. C. Innate Immune Pattern Recognition: A Cell Biological Perspective. Annu. Rev. Immunol. 33, 257–290 (2015).

13. Fitzgerald, K. A. & Kagan, J. C. Toll-like Receptors and the Control of Immunity. Cell 180, 1044–1066 (2020).

14. Reyes Ruiz, V. M., et al. Broad detection of bacterial type III secretion system and flagellin proteins by the human NAIP/NLRC4 inflammasome. Proc. Natl. Acad. Sci. 114, 13242–13247 (2017).

15. Tenthorey, J. L. et al. The structural basis of flagellin detection by NAIP5: A strategy to limit pathogen immune evasion. Science 358, 888–893 (2017).

16. Matico, R. E. et al. Structural basis of the human NAIP/NLRC4 inflammasome assembly and pathogen sensing. Nat. Struct. Mol. Biol. 31, 82–91 (2024).

17. Bobonis, J. et al. Bacterial retrons encode phage-defending tripartite toxin–antitoxin systems. Nature 609, 144–150 (2022).

18. Maffei, E. et al. Systematic exploration of Escherichia coli phage–host interactions with the BASEL phage collection. PLOS Biol. 19, e3001424 (2021).

19. Kim, D. E., et al. De novo design of small beta barrel proteins. Proc. Natl. Acad. Sci. 120, e2207974120 (2023).

20. Tamman, H. et al. A nucleotide-switch mechanism mediates opposing catalytic activities of Rel enzymes. Nat. Chem. Biol. 16, 834–840 (2020).

21. Steinchen, W. et al. Catalytic mechanism and allosteric regulation of an oligomeric (p)ppGpp synthetase by an alarmone. Proc. Natl. Acad. Sci. U. S. A. 112, 13348–13353 (2015).

22. Rousset, F. et al. Phages and their satellites encode hotspots of antiviral systems. Cell Host Microbe 30, 740–753.e5 (2022).

23. Deep, A., Liang, Q., Enustun, E., Pogliano, J. & Corbett, K. D. Architecture and Infection-Sensing Mechanism of the Bacterial PARIS Defense System. http://biorxiv.org/lookup/doi/10.1101/2024.01.02.573835 (2024) doi:10.1101/2024.01.02.573835.

24. Burman, N., et al. Viral Proteins Activate PARIS-Mediated tRNA Degradation and Viral tRNAs Rescue Infection. http://biorxiv.org/lookup/doi/10.1101/2024.01.02.573894 (2024) doi:10.1101/2024.01.02.573894.

25. Tenthorey, J. L., Emerman, M. & Malik, H. S. Evolutionary Landscapes of Host-Virus Arms Races. Annu. Rev. Immunol. 40, 271–294 (2022).

26. Ashkenazy, H. et al. ConSurf 2016: an improved methodology to estimate and visualize evolutionary conservation in macromolecules. Nucleic Acids Res. 44, W344–W350 (2016).

27. Wilkinson, M., Chaban, Y. & Wigley, D. B. Mechanism for nuclease regulation in RecBCD. eLife 5, e18227 (2016).

28. Manalastas-Cantos, K., et al. *ATSAS 3.0* : expanded functionality and new tools for small-angle scattering data analysis. J. Appl. Crystallogr. 54, 343–355 (2021).

29. Duong, M. M., Carmody, C. M., Ma, Q., Peters, J. E. & Nugen, S. R. Optimization of T4 phage engineering via CRISPR/Cas9. Sci. Rep. 10, 18229 (2020).

30. Srikant, S., Guegler, C. K. & Laub, M. T. The evolution of a counter-defense mechanism in a virus constrains its host range. eLife 11, e79549 (2022).

31. Kabsch, W. *XDS*. Acta Crystallogr. D Biol. Crystallogr. 66, 125–132 (2010).

32. McCoy, A. J. et al. *Phaser* crystallographic software. J. Appl. Crystallogr. 40, 658–674 (2007).

33. Mirdita, M. et al. ColabFold: making protein folding accessible to all. Nat. Methods 19, 679–682 (2022).

34. Liebschner, D. et al. Macromolecular structure determination using X-rays, neutrons and electrons: recent developments in *Phenix*. Acta Crystallogr. Sect. Struct. Biol. 75, 861–877 (2019).

35. Emsley, P., Lohkamp, B., Scott, W. G. & Cowtan, K. Features and development of *Coot*. Acta Crystallogr. D Biol. Crystallogr. 66, 486–501 (2010).

36. Cowtan, K. The Buccaneer software for automated model building. 1. Tracing protein chains. Acta Crystallogr. D Biol. Crystallogr. 62, 1002–1011 (2006).

37. Afonine, P. V. et al. Towards automated crystallographic structure refinement with phenix.refine. Acta Crystallogr. D Biol. Crystallogr. 68, 352–367 (2012).

38. Smart, O. S. et al. Exploiting structure similarity in refinement: automated NCS and target-structure restraints in BUSTER. Acta Crystallogr. D Biol. Crystallogr. 68, 368–380 (2012).

39. AltschuP, S. F., Gish, W., Miller, W., Myers, E. W. & Lipman, D. J. Basic Local Alignment Search Tool. 8.

40. Edgar, R. C. MUSCLE: multiple sequence alignment with high accuracy and high throughput. Nucleic Acids Res. 32, 1792–1797 (2004).

41. Perez-Riverol, Y. et al. The PRIDE database resources in 2022: a hub for mass spectrometry-based proteomics evidences. Nucleic Acids Res. 50, D543–D552 (2022).

